# Entrenchment of germline amino-acid differences in antibody affinity maturation

**DOI:** 10.64898/2026.04.21.720000

**Authors:** Noam Harel, Kevin Sung, Will Dumm, Mackenzie M. Johnson, David Rich, Julia Fukuyama, Hugh K. Haddox, Frederick A. Matsen

## Abstract

Entrenchment — epistasis that locks in amino acid differences between homologous proteins, so each disfavors substitutions toward the other’s state — has been demonstrated along individual protein lineages over deep evolutionary time. Antibodies offer a unique system for studying entrenchment: multiple homologous germline V gene paralogs provide diverse starting points, and the rapid somatic evolution of affinity maturation generates dense phylogenies from which selection on germline-encoded residues can be inferred. Using DASM, a deep-learning model that separates selection from mutation in antibody repertoire data, we test for entrenchment across immunoglobulin heavy chain variable (IGHV) genes. We detect entrenchment at two levels of germline divergence, driven by different sources of epistasis. Within V gene families (up to *∼*20% amino acid divergence), entrenched sites cluster at the borders of the complementarity-determining regions (CDRs, the antigen-binding loops) and show high germline diversity. These sites contact antigen, light chain, and the heavy-chain CDR3 loop, all of which vary independently of the IGHV germline. This pattern is consistent with epistasis from genetically uncoupled partners. Between V gene families, at deeper levels of divergence (25–40%), entrenchment additionally includes positions in the framework scaffold distant from binding interfaces, suggesting a larger contribution from intra-heavy-chain structural constraints. Observed mutation frequencies in human repertoires corroborate these predictions where data are sufficient. Together, these results demonstrate that the rapid somatic evolution of antibodies can serve as a lens for revealing epistatic constraints acting on germline-encoded residues, including constraints imposed by genetically uncoupled partners assembled during B cell development.

## Introduction

Amino acid differences between homologous proteins can become entrenched: maintained by epistatic constraints such that each homolog disfavors substitutions toward the other’s state [1]. Previous studies have attributed entrenchment to intra-protein epistasis accumulating along single lineages over deep evolutionary time. For example, experimental studies found that restrictive mutations in steroid hormone receptors created an “epistatic ratchet” preventing reversal [2], and deep mutational scanning of Hsp90 across reconstructed ancestors showed that mutational effects shift pervasively over a billion years of evolution [3]. Consistent with these observations, computational studies showed that entrenchment can arise under purifying selection, as substitutions become contingent on their predecessors and increasingly costly to revert [1, 4]. Beyond these single-lineage comparisons, deep mutational scanning of HIV Env variants was used to quantify levels of entrenchment between two substantially diverged co-existing homologs [5]. Such horizontal comparisons can detect not only classical entrenchment but also other epistatic mechanisms that produce reciprocal incompatibilities; we use “entrenchment” as a functional definition throughout, referring to any reciprocal lock-in regardless of the evolutionary path that produced it.

Antibodies offer an unusually powerful system for studying entrenchment at multiple evolutionary scales. Each human haplotype encodes approximately 50 functional immunoglobulin heavy chain variable (IGHV) genes at a single locus, all paralogs of a common ancestor [6]. These paralogs are conventionally grouped by sequence similarity into seven families (IGHV1–IGHV7) [7], and have diversified along the germline to now differ at up to *∼*40% of amino acid positions (Figure 1A, Supplementary Figure S1). During affinity maturation, B cells expressing these genes undergo rapid somatic evolution inside germinal centers, where activation-induced cytidine deaminase introduces point mutations at rates far above the germline substitution rate [8] and selection for antigen binding acts over days to weeks [9]. This accelerated regime generates densely sampled phylogenies of variants within a single host [10], allowing us to detect entrenchment by inferring selection on germline-encoded residues.

**Figure 1:**
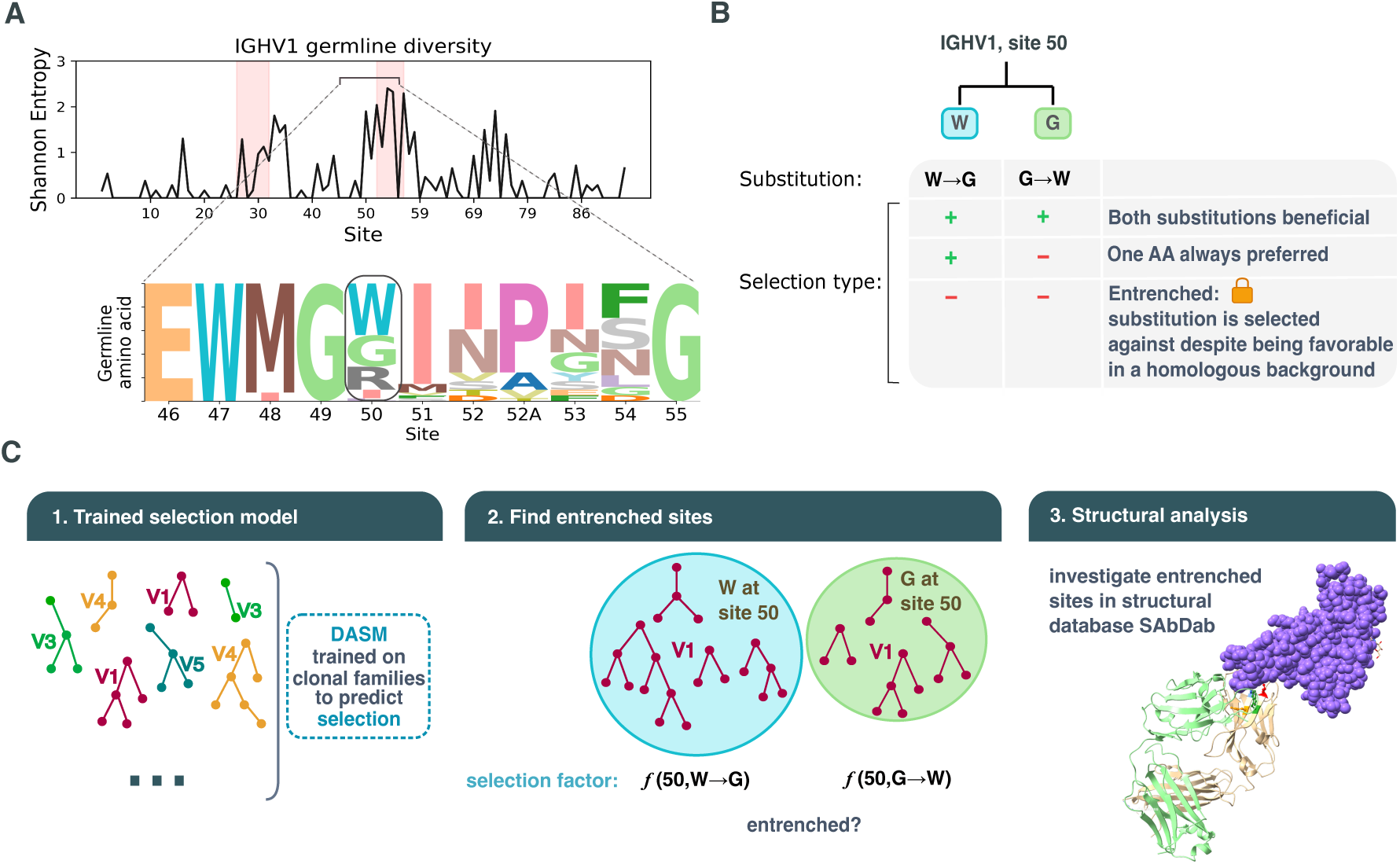
Detecting entrenchment at germline divergent sites using DASM selection factors. **(A)** When V genes have different germline amino acids at homologous positions, epistatic interactions can cause substitutions toward the other V gene’s germline state to be strongly selected against in each genetic background. Examining entropy across sites reveals regions with germline diversity; entrenchment analysis requires such diversity at a site. We select site 50, which has germline diversity, as an example for this figure; in this example it contains a tryptophan (W), a glycine (G), an arginine (R) or other less common options, depending on the V gene. **(B)** To test entrenchment between every two options, for example tryptophan and glycine, we compare the selection on a germline W mutating to G versus a germline G mutating to W. Three outcomes are possible: diversifying selection at the site (substitutions in both directions are favored), directional selection (one amino acid is always preferred), or entrenchment, where both reciprocal substitutions are disfavored, indicating the site resists change toward its homologous state. **(C)** Approach: we use a DASM model trained on clonal families across all V family types to predict selection factors per site, running inference on a held-out test dataset. Sequences are divided by their germline amino acid at the site of interest, if unmutated at that site. We then examine what DASM predicts as the selection factor at the site: in this site 50 example for W*→*G in one group and G*→*W in the other.

Prior work on entrenchment has focused predominantly on intra-protein epistasis as its source. Indeed, Starr et al. [3] found that intra-protein epistasis accounts for nearly all entrenchment in Hsp90, but they noted that the balance between intra- and inter-molecular contributions may differ across protein systems. Antibodies offer a distinctive system: during B cell development, a germline IGHV gene is joined to D and J gene segments by V(D)J recombination, the resulting heavy chain pairs with a light chain from a separate locus, and the assembled complex then binds antigen. During affinity maturation, selection acts in the context of the full complex, so IGHV-encoded residues can experience epistasis from multiple sources: from the IGHV residues themselves, from the D and J gene segments recombined with the IGHV gene, and from the additional proteins the heavy chain interacts with: the light chain and the antigen. Notably, the D and J genes, the light chain, and the antigen are all genetically independent from the IGHV gene and evolve separately, so their constraints would not have been gradually acquired along the germline lineage. Here, we study entrenchment across IGHV genes, asking which germline amino acid differences are entrenched and whether their entrenchment arises from constraints within the IGHV or from interactions with these genetically uncoupled partners.

Beyond identifying the sources of epistasis, entrenchment analysis can reveal which germline differences are selectively maintained rather than neutral [11]. IGHV germline diversity influences V-J pairing [12], heavy-light chain pairing [13, 14], and convergent antibody responses [15]. Certain germline-encoded motifs even bind specific amino acid residues in antigens, driving immunodominant public responses [16]. However, not all germline diversity is necessarily functional: some amino acid differences may be selectively neutral, having fixed through genetic drift.

We test for entrenchment using DASM (Deep Amino acid Selection Model), a deep-learning model trained to quantify selection on germline-encoded amino acids from antibody repertoire data [17]. The model infers selective pressures along phylogenetic trees of millions of affinity-matured antibody sequences. We examine entrenchment at two levels of sequence divergence: within V gene families, where most gene pairs differ at up to *∼*10% for IGHV4, *∼*15% for IGHV1, or *∼*20% for IGHV3, and between families, where they differ at 25–40% (Supplementary Figure S1). Within V gene families, we find that entrenched sites concentrate at the borders of the complementarity-determining regions (CDRs, the antigen-binding loops) and contact multiple genetically independent partners, and we characterize these interactions using structural and physicochemical analyses. Between V gene families, additional entrenched sites appear in framework regions (the scaffolding positions between CDRs), and we examine candidate structural explanations at selected sites. Validation using observed mutation frequencies in human repertoires corroborates these predictions at sites with sufficient data. Together, these results suggest that entrenchment in antibodies arises from both intra-heavy-chain structural constraints and epistasis with genetically uncoupled partners.

## Results

### Identifying entrenched germline amino acids within V families

Under entrenchment, reciprocal substitutions between two germline amino acids found at the same site (e.g., W*→*G and G*→*W) are strongly disfavored (Figure 1B). DASM selection factors quantify how selection shifts the probability of each substitution relative to neutral expectation, given the parent amino acid sequence: a log selection factor above zero indicates the substitution has a beneficial effect, a value of zero indicates a neutral effect, and a value below zero indicates a deleterious effect. We classify a site as entrenched for a given amino acid pair when both reciprocal substitutions have natural-log selection factors below *−*1, i.e., occurring at less than 37% of the neutral rate (Figure 1). Because the strength of entrenchment is continuous, we chose this cutoff to focus on sites with the strongest predicted effects. Under this criterion, a site can be entrenched only if its germline amino acids differ across V genes, and selection during affinity maturation disfavors the reciprocal substitutions between them. The two directions are measured in different clonal families: we estimate selection on *A → B* only in families whose germline carries *A* at the site, and on *B → A* only in those carrying *B*.

We began by examining entrenchment between V genes within the same V family, where most gene pairs differ at less than *∼*20% of their amino acids (Supplementary Figure S1). To detect entrenchment, we identified sites where different V genes have divergent germline amino acids and used DASM to compare predicted selection factors for reciprocal substitutions between these residues (Figure 1). We applied DASM to a repertoire dataset held out from the model training data, restricting to sequences at the most recent common ancestor of each clonal family, where training data are most abundant and selection factors are not yet shifted by somatic mutations accumulated during affinity maturation. We further restricted to single-nucleotide substitutions, as multi-nucleotide substitutions are rare in the training data and may yield less reliable selection-factor estimates.

For each site and pair of germline amino acids, we measured the median predicted log selection factor for substitutions converting one amino acid toward the other. This approach aggregates across all V genes within a family that share the same amino acid at a given site, yielding more robust median log selection factor estimates. Per-V-gene-allele medians confirmed the consistency of this grouping: the intraclass correlation coefficient (ICC) was at least 0.96 for every family (Supplementary Figure S2, Methods).

### Entrenchment between V genes within the same V family clusters at CDR borders

We focused our within-family entrenchment analysis on IGHV1 and IGHV3, the two families with the greatest germline diversity and abundant repertoire data. Across all sites, we tested for entrenchment among 95 and 130 pairs of germline amino acids in IGHV1 and IGHV3, and found 8 and 11 entrenched, respectively, spanning 3 and 5 sites (Figure 2A). Most entrenched sites are retained under stricter thresholds down to *−*1.4 (Supplementary Figures S3, S4, and S5).

**Figure 2:**
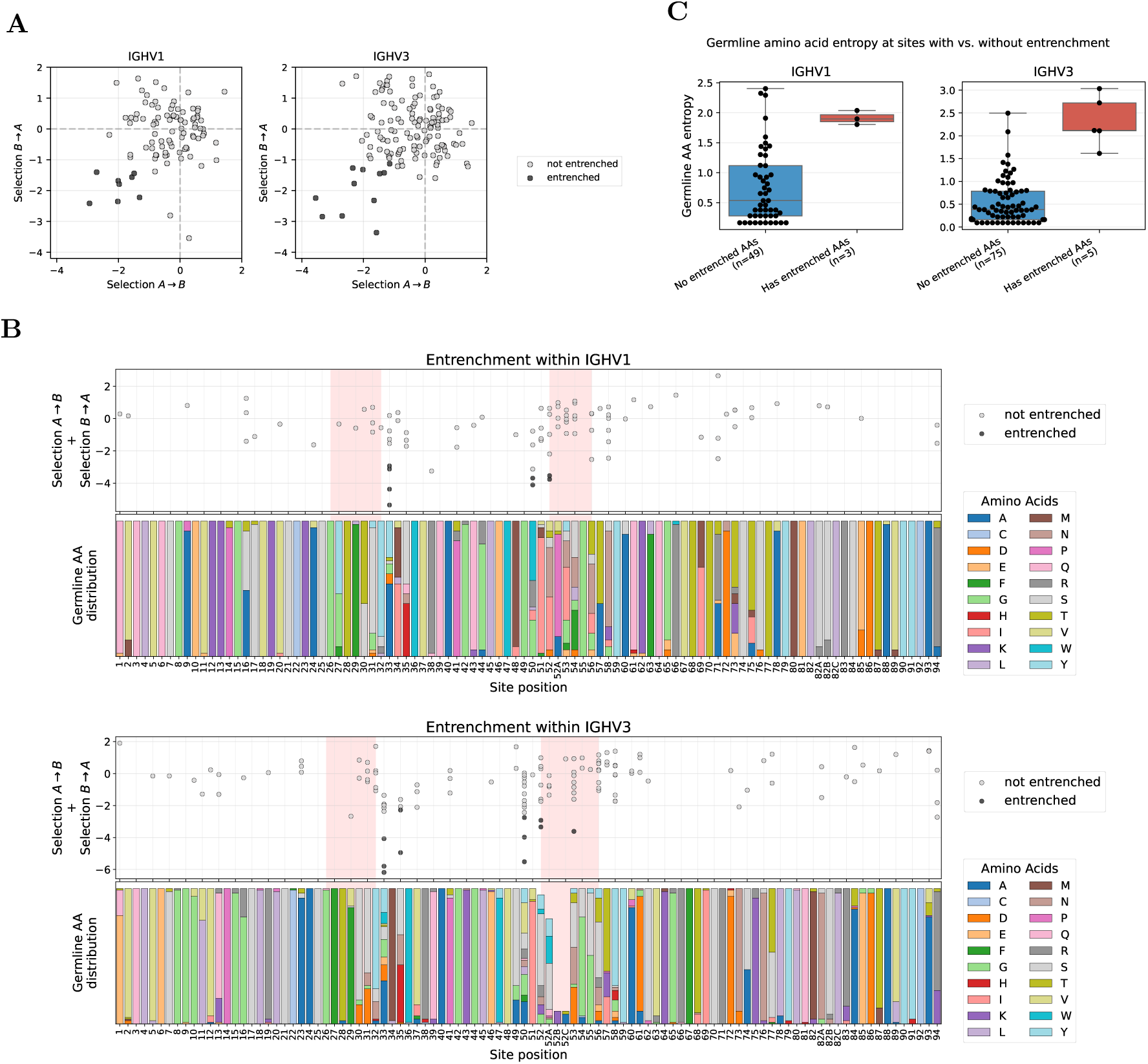
Entrenchment analysis within V families identifies sites at CDR borders with high germline diversity. **(A)** Reciprocal median log selection factors between amino acids A and B (selection *A → B* and selection *B → A*; see Methods), for amino acid pairs at sites where V genes within a family differ in germline identity, shown for IGHV1 (left) and IGHV3 (right). Selection factors here and throughout refer to selection during affinity maturation. Each dot represents a combination of site and amino acid pair; entrenched pairs (both reciprocal median log selection factors *< −*1) are highlighted as black dots. **(B)** Entrenchment mapped along the V gene sequence (IGHV1 top, IGHV3 bottom). Top subpanels show affinity-maturation selection: the sum of reciprocal median log selection factors (selection *A → B* + selection *B → A*) for each amino acid pair at each site, with entrenched pairs highlighted as in (A). Pink shading highlights CDR regions. Bottom subpanels show germline-level diversity: the distribution of germline amino acids across V genes in each family. Sites with germline variability but no points (e.g., site 2 in IGHV3) lack sufficient data after filtering for single-nucleotide substitutions and requiring *≥*10 observations per amino acid (see Methods for full filtering criteria). Analogous analysis for IGHV4 is shown in Supplementary Figure S6. **(C)** Shannon entropy of the germline amino acid distribution at sites with versus without entrenched pairs. Sites with entrenched pairs show higher germline diversity, though there is overlap between distributions. Sites with only one germline amino acid across V genes in the family (entropy = 0) are excluded. A complete list of all entrenched pairs is provided in the project GitHub repository. Site numbering follows the Chothia scheme [18] throughout.

Sites showing entrenchment (hereafter, entrenched sites) cluster predominantly at CDR boundaries (Figure 2B). Specifically, these sites concentrate at the end of CDR1 and beginning of CDR2 in both V families, a pattern also observed in IGHV4 (Supplementary Figure S6). We also detect a single entrenched amino acid pair at CDR2 site 53 in IGHV3 (usually not adjacent to site 52 due to Chothia insertion codes such as 52A). These sites exhibit high germline diversity (Figure 2B, bottom panels). Because entrenchment requires at least two germline amino acids at a site, entrenched sites necessarily have nonzero diversity; however, high diversity alone does not predict entrenchment. The Shannon entropy distributions of entrenched and non-entrenched sites overlap (Figure 2C), and several highly diverse sites show no evidence of entrenchment (e.g., sites 52A–54 in IGHV1 and 52A, 56–58 in IGHV3; Figure 2B).

### Entrenched sites contact multiple genetically independent partners

We next asked what molecular interactions might cause the entrenchment. When we mapped the entrenched sites onto heavy-chain structures, we found a striking pattern: four of the five sites (33, 35, 50, and 52) cluster closely together on two strands of a beta sheet at the base of the CDR1 and CDR2 loops, right at the interface of the heavy-light-antigen complex (Figure 3A–C). These sites often contact three regions whose sequences are genetically independent of the IGHV germline: the light chain (encoded at a separate locus), the antigen, and the largely independent heavy-chain CDR3 loop (CDR-H3; encoded by D and J gene segments and the very end of the V gene, joined by V(D)J recombination with additional junctional diversity).

**Figure 3:**
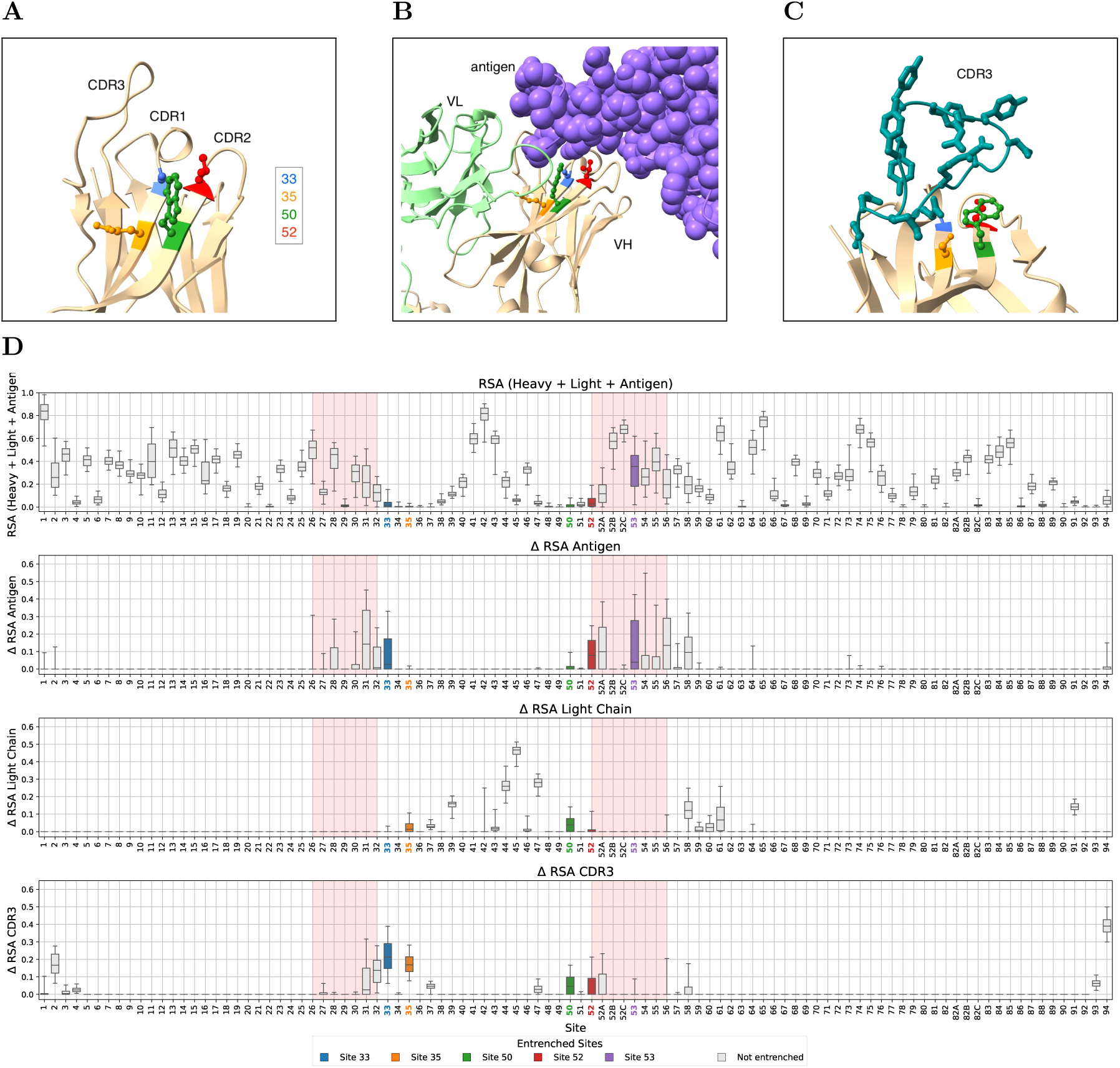
Within-family entrenched sites lie on CDR-scaffolding beta sheets and contact genetically independent partners. **(A)** Heavy chain of an example antibody (PDB: 7L7D, resolution 2.5Å), an IGHV1 antibody binding the SARS-CoV-2 spike protein, showing the spatial arrangement of entrenched sites 33, 35, 50 and 52 with annotated CDRs. These sites cluster at or near the edges of beta sheets that scaffold the CDR loops. **(B)** Same structure with light chain (green) and antigen (purple). Entrenched sites contact the antigen and light chain. **(C)** Heavy chain of a second IGHV1 antibody (PDB: 6B0G, resolution 1.9Å), an IGHV1-18 antibody binding Pfs25, showing the CDR-H3 loop (heavy-chain CDR3, teal) contact with entrenched sites. In this structure, these sites contact the CDR-H3 but not antigen or light chain, contrasting with (B). **(D)** RSA analysis for IGHV3 family, with within-family entrenched sites highlighted. Row 1: Relative solvent accessibility (RSA) in the full complex. Row 2: Change in RSA upon antigen removal (antigen effect). Row 3: Change in RSA upon light-chain removal (light-chain effect). Row 4: Change in RSA upon CDR-H3 removal (positions 95–102). The corresponding RSA analysis for IGHV1 family is shown in Supplementary Figure S7.

To systematically quantify these interactions, we examined antibody structures from the Structural Antibody Database (SAbDab) that include a heavy chain, light chain, and antigen [19] (733 structures for IGHV1 and 1047 for IGHV3; see Methods for filtering criteria). For each structure, we computed the relative solvent accessibility (RSA) of each heavy-chain residue in the full complex, as well as the change in RSA (ΔRSA) upon removing the antigen, the light chain, or the CDR-H3. Throughout, we take the CDR-H3 to span positions 95–102, one position beyond the Chothia boundaries [18] at each end, so that the variable part of the V(D)J junction is included. While the IGHV gene templates part of the first codon or two of this region, the number of nucleotides trimmed during V(D)J recombination varies from rearrangement to rearrangement [20, 21], so the amino acids at its start differ among antibodies carrying the same V allele. Unlike the CDR-H3 loop, the adjacent framework region 4 (FR4) is a conserved framework segment that presents consistent selection pressure across antibodies, so it was retained despite also being J-gene-encoded. The four clustered entrenched sites (33, 35, 50, 52) are highly buried in the full complex, while site 53 is more solvent exposed (Figure 3D, top row). All three genetically independent partners contribute to burial at these sites, with the relative importance varying by position: sites 33 and 52 are buried primarily by antigen and the CDR-H3 loop, site 35 by the CDR-H3 loop and light chain, site 50 roughly equally by all three, and site 53 primarily by antigen (Figure 3D, rows 2–4).

To further characterize these interactions, we examined both the physicochemical properties and the structural contacts of amino acid pairs at entrenched sites. Entrenched amino acid substitutions involve more physicochemically distinct amino acids than non-entrenched substitutions at the same sites, as measured by Grantham distance (stratified permutation test across 8 V-family *×* site groups, *p <* 0.0001; Figure 4). Moreover, different germline amino acids at the same site establish distinct contact patterns with antigen, light chain, and CDR-H3 (Supplementary Figure S8), though some variability exists among structures with the same amino acid. At site 33, where amino acid pairs are highly repeated between IGHV1 and IGHV3, most of the same amino acid pairs are entrenched in both families (e.g., aspartic acid (D)*↔*glycine (G) and aspartic acid (D)*↔*tyrosine (Y)), while other, more physicochemically similar pairs are tolerable in both (e.g., alanine (A)*↔*glycine (G) and serine (S)*↔*threonine (T)), consistent with the same Grantham distance pattern (Figure 4).

**Figure 4:**
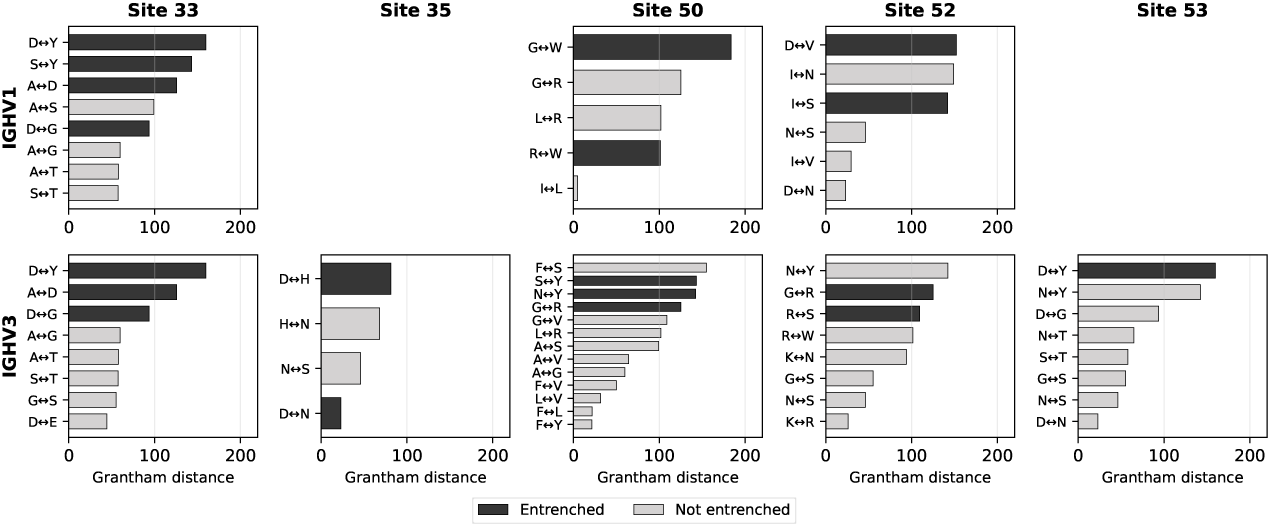
Entrenched amino acid pairs are more physicochemically distinct than the other pairs at their sites. Grantham distance for each amino acid pair at the five within-family entrenched sites, shown separately for IGHV1 (top row) and IGHV3 (bottom row). Sites 35 and 53 appear only in IGHV3, as they are not entrenched within IGHV1. Pairs are sorted by Grantham distance; black bars indicate entrenched pairs and grey bars indicate non-entrenched pairs. Within each site, entrenched pairs tend to have larger Grantham distances, indicating greater physicochemical difference between the two amino acids. A stratified permutation test across 8 V-family *×* site groups confirms this trend (*p <* 0.0001; 10,000 permutations).

Together, these results suggest that entrenchment of V-gene residues could arise through epistatic interactions between the V gene and each of these binding partners. The high within-family germline diversity at these sites indicates that the constraints are not imposed by the heavy-chain fold alone, since closely related V genes tolerate diverse amino acids at these positions. However, not all sites with partner contact are entrenched: sites 31 and 32 have both antigen and CDR-H3 loop contact without being entrenched, and multiple other sites have contact with one partner without being entrenched (Figure 3D), suggesting that partner contact alone is not sufficient.

### Entrenchment between V families suggests intra-protein structural constraints

We next examined entrenchment between more distantly related proteins by comparing V genes from different families. V genes from different families typically differ in approximately 25–40% of their amino acids (Supplementary Figure S1), representing evolutionary divergence that occurred over a longer timescale than within-family variation. We applied a similar approach to identify entrenched sites between V families, focusing on sites with germline divergence and assessing reciprocal purifying selection patterns. Again, groups of V genes within a V family that share the same amino acid at a site were aggregated to obtain more robust median log selection factor estimates for that site.

The between-family analysis predicted entrenchment at several sites, including four of the five sites predicted to be entrenched in the within-family analysis (Figure 5). Sites that were only predicted to be entrenched in the between-family analysis occur predominantly in framework regions and span a range of solvent accessibility values, from buried to highly exposed (Figure 5C, Supplementary Figure S9). These sites show less within-family germline diversity than within-family entrenched sites (Supplementary Figure S10). Several, such as sites 9 and 73, are among the most solvent-exposed positions in the heavy chain. Most of these sites do not make contact with antigen, light chain, or CDR-H3, though site 94, which borders the CDR-H3 loop, contacts both it and the antigen (Supplementary Figure S9). For the majority of between-family-only entrenched sites, the lack of contact with genetically independent partners suggests that their entrenchment arises from structural constraints intrinsic to the part of the heavy chain encoded by the V gene rather than from epistasis with genetically uncoupled partners.

**Figure 5:**
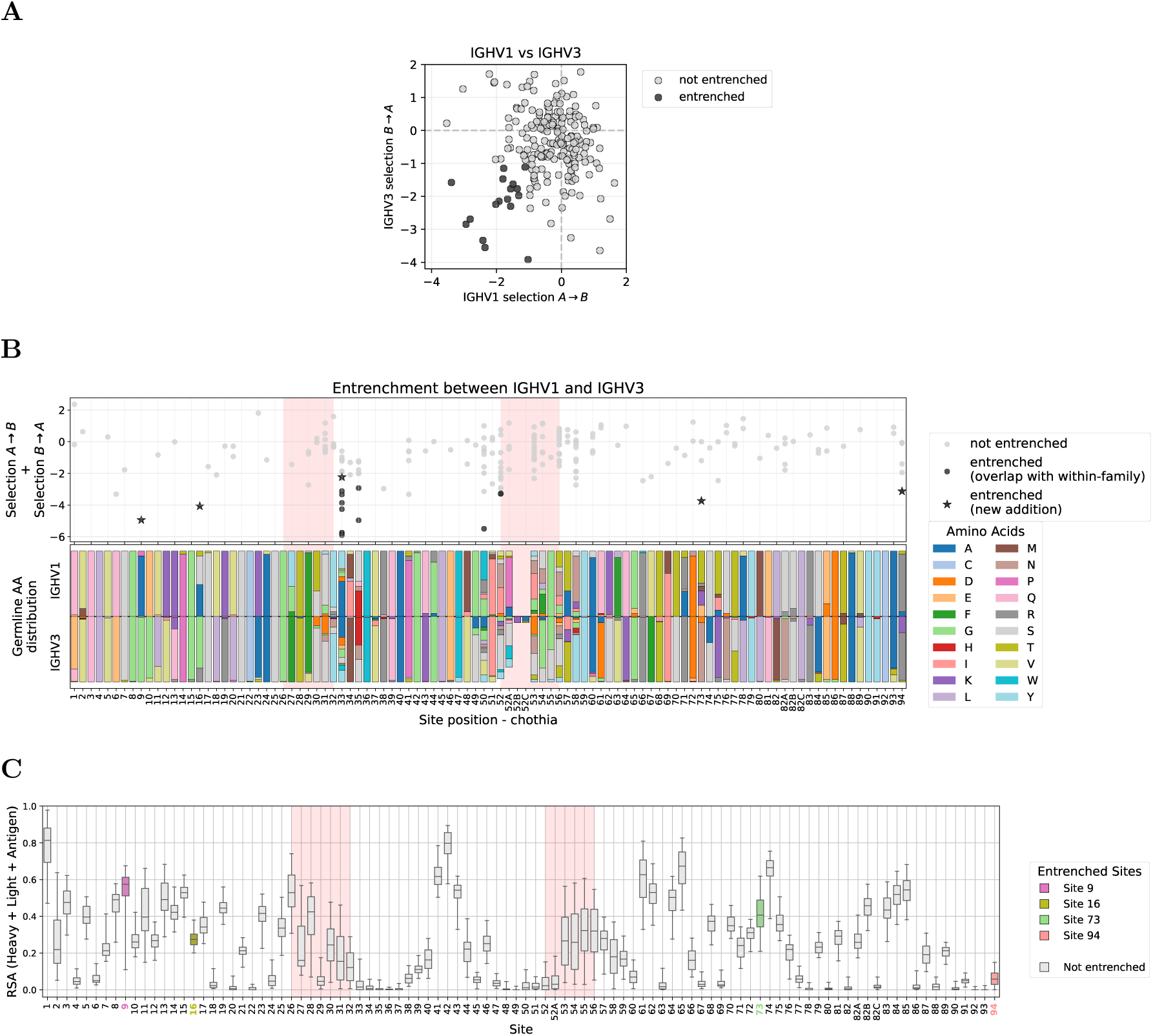
Entrenchment between more distantly related homologs retains within-family sites and adds framework positions. **(A)** Reciprocal median log selection factors (selection *A → B* vs. selection *B → A*) for amino acid pairs at sites where IGHV1 and IGHV3 differ in germline identity; entrenched pairs (both *< −*1) are highlighted. **(B)** Entrenchment between IGHV1 and IGHV3 mapped along the V gene sequence. Top panel shows the sum of reciprocal median log selection factors for each amino acid pair at each site. Light grey dots indicate non-entrenched pairs; dark dots indicate entrenched pairs that overlap with within-family entrenched sites; stars indicate entrenched pairs not entrenched within either family (new additions). Pink shading highlights CDR regions. Bottom panel shows the distribution of germline amino acids at each site in IGHV1 (above axis) and IGHV3 (below axis). Analogous analysis for IGHV3 vs IGHV4 and IGHV1 vs IGHV4 is shown in Supplementary Figure S11. **(C)** Relative solvent accessibility (RSA) in the full antibody-antigen complex for IGHV1, highlighting sites entrenched only in the between-family analysis (i.e., not entrenched within either family). Extended RSA analysis including antigen, light chain, and CDR-H3 effects for both IGHV1 and IGHV3 is shown in Supplementary Figure S9. These between-family-only sites show lower germline diversity than within-family entrenched sites (Supplementary Figure S10).

We investigated several of these entrenched positions in detail and found that they lie in regions with structural differences between IGHV1 and IGHV3; however, it is not immediately obvious how these differences give rise to entrenchment. For example, site 9 has an entrenched alanine/glycine polymorphism between IGHV1 and IGHV3 and lies in a region spanning sites 7–10 where the two families adopt distinct backbone conformations, in contrast to the generally conserved backbone geometry in most of the remaining structure (Figure 6A; Supplementary Figure S12). The distinct backbone conformations at this region between IGHV1 and IGHV3 are consistent with entrenchment at this site, though the precise mechanism by which these structural differences constrain amino acid identity remains unclear. Site 73 has an asparagine/lysine polymorphism between IGHV3 and IGHV1 and lies in a loop spanning sites 72–75. In IGHV3, asparagine at site 73 is consistently anchored by hydrogen bonds to arginine at site 71 and a CDR-H2 backbone carbonyl across all examined structures, while lysine at site 73 in IGHV1 forms no framework hydrogen bonds and only inconsistent contact with CDR-H2 (Figure 6B–C, Supplementary Table S1). This provides a partial structural explanation: replacing asparagine with lysine in IGHV3 would disrupt these stabilizing contacts, though the structural basis for why the reverse substitution would be disfavored in IGHV1 is less clear.

**Figure 6:**
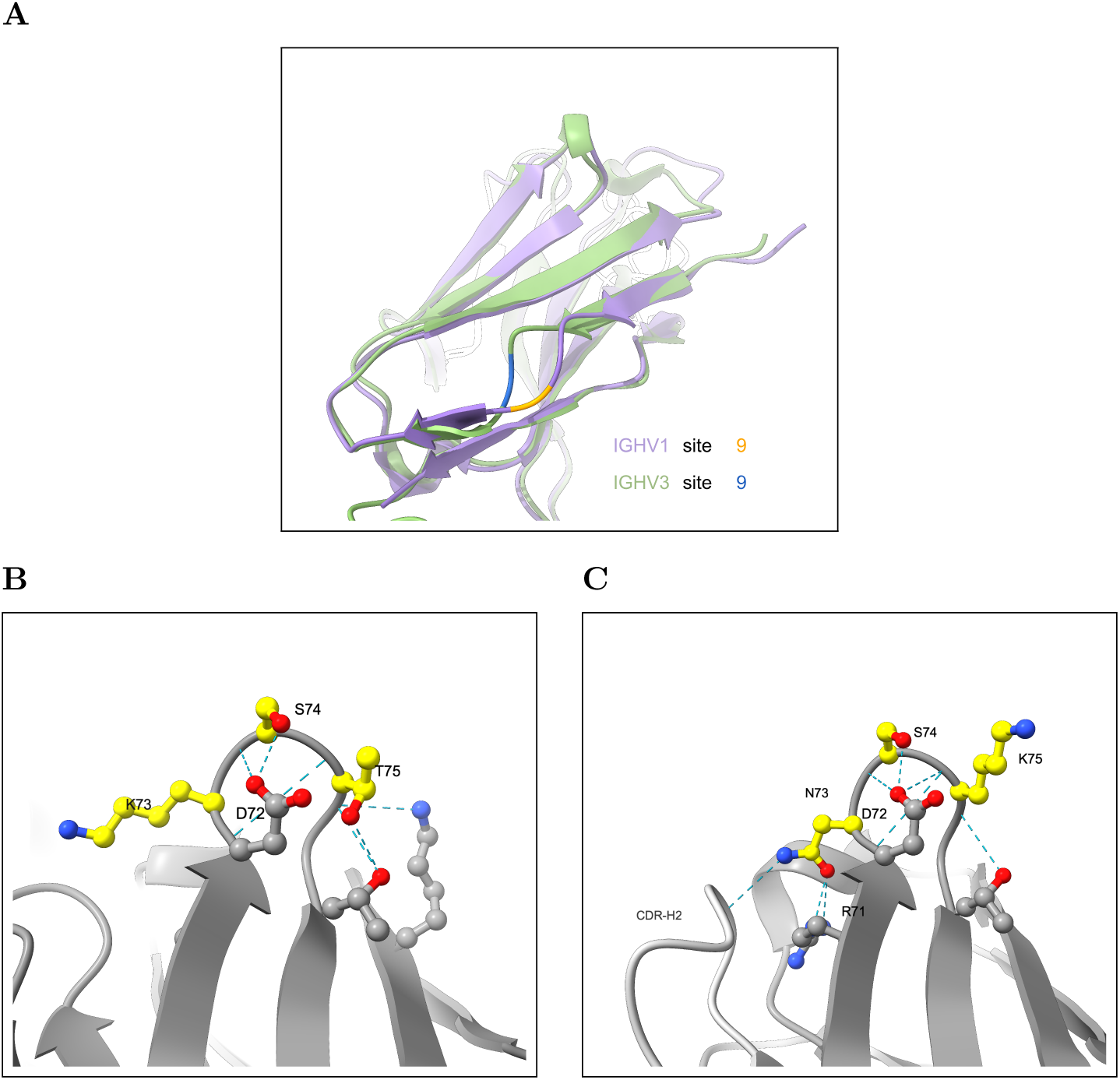
Entrenched sites 9 and 73 are in regions with structural differences between IGHV1 and IGHV3. **(A)** Structural alignment of IGHV1 (purple; PDB: 7VYT, 1.53 Å resolution) and IGHV3 (green; PDB: 8FFE, 1.72 Å resolution) near site 9, where IGHV1 carries alanine (orange) and IGHV3 carries glycine (blue). The two families adopt divergent backbone conformations at sites 7–10. Backbone dihedral angle analysis across 521 IGHV1 and 962 IGHV3 SAbDab structures is shown in Supplementary Figure S12. **(B)** IGHV1 hydrogen bonding network (PDB: 2NY1): K73 lacks consistent framework hydrogen bond partners, while T75 anchors inward via hydrogen bonds. Sites 73–75 are highlighted in yellow and hydrogen bonds are shown as cyan dashes. **(C)** IGHV3 hydrogen bonding network (PDB: 4H8W): N73 is anchored by hydrogen bonds to R71 and a CDR-H2 backbone carbonyl; colored as in (B). H-bond analysis across all eight examined structures is reported in Supplementary Table S1. Density supporting the side chain placements in (B) and (C) is shown in Supplementary Figure S13.

### Validation of DASM predictions using mutation rate ratios

DASM is a transformer-encoder model that quantifies how selection shifts the probability of each substitution relative to neutral expectation [17]. Neutral expectations are provided by Thrifty, a convolutional neural network over learned 3-mer embeddings that estimates the probability of each nucleotide substitution from local sequence context [22]. To validate these predictions, we independently estimated selection using a simplified approach based on mutation counts.

Purifying selection means fewer changes than somatic hypermutation alone would produce, not low diversity in absolute terms, so this approach compares observed mutation counts to a neutral expectation. To do so, we calculated the ratio of observed to expected mutation rates for each site and substitution by comparing mutation counts in productive sequences, which are subject to selection, to counts in out-of-frame (non-productive) sequences, which evolve neutrally with respect to amino acid changes [23, 22], accounting for phylogenetic branch lengths. For the productive mutation counts, we used the same held-out repertoire dataset used for entrenchment detection. For the out-of-frame counts, we used a separate dataset of non-productive sequences, which persist in B cells whose functional receptor is encoded by the other allele and are therefore not subject to selection on amino acid changes. This dataset was not used to train the neutral mutation model, though it was used to fit the multi-hit correction (3 parameters only; see Methods). To ensure stable rate ratio estimates, we required *≥* 5 mutation counts in the out-of-frame data, retaining 1,180 out of 3,218 substitutions.

For substitutions passing this filter, the log ratio of observed to expected mutation rates correlated with DASM’s predicted log selection factors (Figure 7A), consistent with the model capturing genuine selective constraints. This approach could directly validate only 21 of the 83 distinct entrenched substitutions (all comparisons, including IGHV4), as many had too few mutations in the out-of-frame sequences for stable rate ratio estimates. Of those 21, all showed purifying selection and 18 (86%) fell below the *−*1 entrenchment threshold, including substitutions at each of the four CDR-bordering sites (33, 35, 50, and 52). Non-productive sequences are far fewer than productive ones, making the out-of-frame data the limiting factor in this analysis.

**Figure 7:**
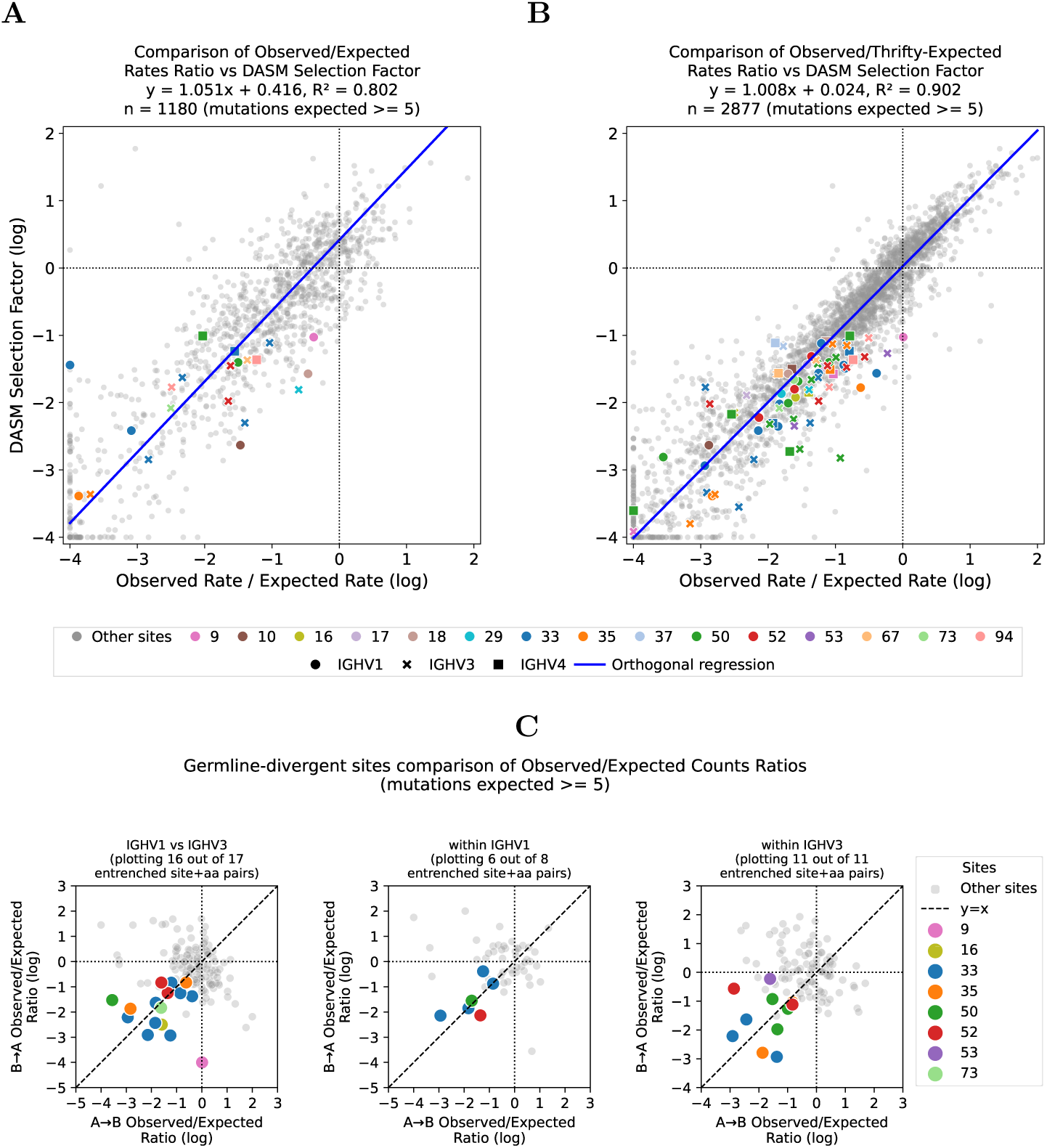
Observed mutation rate ratios support DASM entrenchment predictions where data are sufficient. **(A)** Validation using out-of-frame sequences as neutral baseline. The log ratio of observed to expected mutation rates correlates with median log selection factors predicted by our model for sites and substitutions with sufficient mutation counts (expected counts *≥* 5). Each point represents one V-family, site, parent amino acid, and target amino acid combination. Correlation is calculated for all points; entrenched sites and substitutions that pass filtering are highlighted in color. **(B)** Validation using Thrifty-inferred expected mutation rates. The log ratio of observed mutations (from productive sequences) to expected mutations (inferred from Thrifty neutral model) correlates with median log selection factors predicted by DASM. This approach allows validation of more sites compared to the out-of-frame-based approach, while maintaining strong correlation with DASM predictions. Entrenched sites and substitutions that pass filtering are highlighted in color. **(C)** Pairwise validation of entrenched substitutions. Log rate ratios in both directions (A*→*B and B*→*A) as calculated in (B) for all possible pairs of reciprocal substitutions, while highlighting those identified as entrenched by DASM. Not all entrenched substitutions that appear in (B) appear here, as some do not have sufficient data for both directions of the pair to be included in this plot. Analogous validation for comparisons involving IGHV4 is shown in Supplementary Figure S14.

We thus used a complementary approach: instead of out-of-frame data as neutral baseline, we inferred expected counts directly from Thrifty and used the much larger training dataset to increase statistical power (see Methods). Because Thrifty estimates neutral rates from local sequence context rather than per-site counts, it should provide more stable expected rates at sites where out-of-frame data are too sparse for reliable empirical estimates, though confirming this directly would require larger out-of-frame datasets. This retained 2,877 out of 3,247 substitutions after applying the same filter.

Using this approach, the correlation between DASM selection factors and observed-to-expected mutation rate ratios remained strong (Figure 7B). To validate predicted entrenchment for specific substitution pairs, we examined whether both directions of each reciprocal pair show purifying selection (Figure 7C). Of 36 entrenched reciprocal pairs, 33 had sufficient data in both directions; of these, 29 (88%) showed log rate ratios below *−*0.5 and 19 (58%) below *−*1 in both directions, covering all four CDR-bordering sites. The sole exception to purifying selection was IGHV1 site 9 A*→*G, which had a log rate ratio near zero. This discrepancy arises because the gene IGHV1-18 has atypically positive selection at this site (Supplementary Figure S2) and contributes disproportionately to the expected counts due to higher neutral mutability, so it dominates the aggregate observed-to-expected ratio despite most V genes in this family showing strong purifying selection. The median-based aggregation across V genes in our analysis is robust to such outliers.

All entrenched amino acid substitutions from both within- and between-family comparisons, along with their DASM-based selection factors and mutation rate ratios from both validation approaches, are available in the project repository.

## Discussion

Previous studies have attributed entrenchment to intra-protein epistasis accumulating over deep evolutionary time [2, 3, 1, 4]. Our results suggest that epistasis from genetically uncoupled partners can also drive entrenchment: within V gene families, entrenched sites contact antigen, light chain, and the CDR-H3 loop, and are conserved during affinity maturation even though the same sites tolerate diverse amino acids across the germline repertoire.

The relative contribution of genetically uncoupled and intra-IGHV epistasis shifts with evolutionary distance. Among closely related V genes, intra-chain structural constraints appear to be largely shared; entrenched sites instead cluster at CDR borders, where interactions with antigens, light chains, and CDR-H3 loops could lock in germline amino acids during affinity maturation. At deeper divergence between V gene families, additional entrenched positions appear, which seem likely to be due to intrinsic structural requirements of the heavy-chain region encoded by the V gene, while the CDR border sites identified within families largely persist (Figure 5). In both cases, the constraints predate affinity maturation: V(D)J recombination, heavy-chain folding, light-chain pairing, and antigen binding all occur beforehand, so affinity maturation reveals rather than creates these constraints. This distinguishes our analysis from recent experimental studies that characterize epistasis among somatic mutations during affinity maturation [24, 25]: those studies map how acquired mutations interact as antibodies evolve within a lineage, whereas the constraints we detect are germline-encoded.

Our analysis aggregates selection signals across thousands of clonal families with diverse CDR-H3, light chain, and antigen combinations, averaging out partner-specific constraints. The entrenchment that persists reflects general constraints on IGHV-encoded residues that hold across the combinatorial diversity of partners encountered during affinity maturation. Different V genes carry different amino acids at these same positions, each favored by selection in its own sequence context, suggesting that the entrenched residues help define each V gene’s distinct mode of engaging its partners.

Within-family entrenchment concentrates at CDR1 and CDR2 edges rather than in CDR centers. CDR-central sites tend to be more solvent-exposed, even in the full antibody-antigen complex, and among the genetically independent partners, they primarily contact only the antigen rather than the light chain or CDR-H3 loop. Additionally, many CDR-central sites experience diversifying selection during affinity maturation [26], which works against the reciprocal purifying selection that defines entrenchment. CDR border sites, by contrast, occupy multiple roles: they facilitate contact with the CDR-H3 loop, light chain, and antigen, and they are also part of the framework that scaffolds CDR1 and CDR2. These multiple structural and functional roles might make entrenchment more common at these sites. Entrenched sites partially overlap with germline-encoded antigen-binding (GRAB) motifs [16] at sites 33, 52, and 94, but the two approaches capture different signals: GRAB motifs identify residues that contact specific antigen amino acids, whereas entrenchment detects any site under reciprocal purifying selection, regardless of the mechanism. Conversely, some GRAB sites may not appear entrenched because DASM averages selection across thousands of clonal families with diverse antigens, which could mask antigen-specific constraints.

The antibody system combines properties rarely available together: dozens of homologous germline genes, combinatorial assembly by V(D)J recombination and chain pairing, diverse antigens, and fast somatic evolution [10]. This environment offers a unique opportunity to observe multiple levels of epistatic constraints: within a single germline-encoded segment (intra-IGHV constraints), between independently recombined segments of the same polypeptide (IGHV versus CDR-H3), and between separate proteins that function as a complex (heavy chain versus light chain and antigen). Because each B cell assembles a new combination of these before affinity maturation, the system is a natural experiment for detecting epistasis from genetically uncoupled regions.

Because entrenched residues appear to help define each V gene’s distinct mode of engaging its partners, selection may act to maintain the variety of amino acids at these sites across V genes, which would help explain why the germline carries so many V genes. The major histocompatibility complex (MHC) offers a precedent for selection maintaining diversity among immune genes: an individual carries at most two alleles at each of a few loci, so diversity persists at the population level, where balancing selection preserves thousands of alleles [27, 28]. An individual carries roughly 50 IGHV paralogs per haplotype, so diversity could be preserved within a single genome, with selection favoring divergence among paralogs at entrenched sites because loss of any one could narrow the range of CDR-H3 and light-chain combinations the repertoire can accommodate. Testing this would require germline-level analysis across the IGHV locus, long impeded by assembly difficulties [29, 30] but increasingly tractable with population-scale genotyping [7].

Four methodological limitations should be noted. First, we restricted our analysis to single-nucleotide substitutions, so not all potentially entrenched substitutions can be assessed. Second, affinity maturation could itself generate new entrenchment as compensatory somatic mutations accumulate among somatically acquired residues, a possibility our germline-focused framework does not address. Third, not all DASM predictions can be independently validated with counts, as both the out-of-frame and Thrifty-based approaches retain only a subset of testable substitutions. Where both are available, the strong correlation between DASM and counts-based estimates supports the model’s ability to generalize. Model-based inference can extend predictions to data-sparse sites where counts-based approaches lack statistical power, but confirming whether these predictions reflect genuine biological constraints will require larger repertoire datasets or experimental validation. Fourth, closely related V alleles that differ at only a few nucleotides can be confused during germline inference when somatic hypermutation hits a differing site, potentially mislabeling a somatically mutated residue as germline at that site. Our simulations show that such misassignment is negligible at most entrenched sites, but at IGHV4 site 29 we cannot fully distinguish germline-encoded entrenchment from entrenchment of somatic mutations acquired before the clonal family root (see Methods and Supplementary Figure S15).

Epistasis with genetically uncoupled partners predicts that selection at the entrenched sites differs between clonal families with different light chains, antigens, and CDR-H3s. Of these partners, only the CDR-H3 lies within DASM’s input sequence. We can therefore ask whether selection at those sites depends on the CDR-H3, using its length as a simple summary of the loop. At the sites the CDR-H3 contacts, including the within-family entrenched ones, selection shifts more with length than elsewhere in the V region (Supplementary Figure S16). The shift is also directional: for most entrenched substitutions, selection against leaving the germline amino acid is stronger in families with longer CDR-H3s, as the structures predict, since a longer CDR-H3 buries these sites more. The shift is small, as expected, since length compresses the biochemical and conformational variety of a complex region into a single parameter. Recent experimental work offers independent support: Phillips et al. [25] found epistasis between heavy- and light-chain mutations in the CH65 broadly neutralizing anti-influenza antibody, including at heavy-chain sites 33, 35, and 52, which we identify as entrenched within V families. Their combinatorial library, however, varied somatic mutations within a single lineage. Extending deep mutational scanning [31] to germline substitutions across diverse light-chain and CDR-H3 contexts would test directly whether epistasis with genetically uncoupled partners constrains these germline differences.

More broadly, our findings illustrate that constraints from genetically independent partners — whether inter-molecular or inter-segmental — can shape germline-encoded residues in ways that are difficult to detect when proteins are analyzed in isolation. Somatic evolution within the antibody repertoire provides a rare opportunity to observe these constraints directly, and similar approaches may extend to other systems where related proteins evolve together with diverse partners.

## Materials and methods

### Data, code, and models

We used four human heavy-chain antibody repertoire datasets (Table 1). The productive data were processed as in the DNSM (Deep Natural Selection Model) [26] and DASM publications [17], and the out-of-frame data as in the Thrifty publication [22]. Sequences from each dataset are partitioned into clonal families with inferred phylogenetic trees. Each edge of a phylogenetic tree gives a parent– child pair (PCP): a parent sequence and the child sequence descended from it. These PCPs are the unit of analysis throughout.

**Table 1:** Antibody repertoire datasets used in this study. PCP counts reflect all parent–child pairs in the source dataset. Individual analyses in this paper apply additional filters, including restriction to depth=2 parent–child pairs, as described in the relevant Methods subsections.

| Dataset | Sequence type | Indi-viduals | Clonal families | PCPs | Role in this study | Ref. |
|---|---|---|---|---|---|---|
| Jaffe | Productive | 4 | 57,364 | 228,790 | DASM training dataset; here used for Thrifty-based validation of DASM selection factors (Approach 2). | [32] |
| Tang | Productive | 21 | 146,920 | 522,586 | DASM training dataset; here used for Thrifty-based validation of DASM selection factors (Approach 2). | [33, 34] |
| Rodriguez | Productive | 50 | 7,769 | 21,754 | Held-out test dataset (not used for DASM training); used for entrenchment inference from DASM selection factors and for out-of-frame-based validation (Approach 1). | [30] |
| Tang-SHM | Out-of-frame (non-productive) | 21 | 3,209 | 9,984 | Neutral baseline for validation (Approach 1); held-out test dataset for Thrifty, except for its multi-hit correction component (3 parameters) which was fit on these data. | [33, 34] |

Out-of-frame sequences arise when V(D)J recombination introduces a frameshift near the CDR-H3, leaving the rearrangement unable to encode a receptor [23]. A cell retains such a rearrangement whenever its other immunoglobulin allele recombined productively, with allelic exclusion confining expression to that productive allele. The unexpressed copy is nonetheless carried through the germinal center and undergoes somatic hypermutation in parallel with its expressed counterpart, so its mutations accumulate free of amino acid-level selection [23, 22].

The PCP counts for Jaffe and Tang in Table 1, which are used for validation (Approach 2) of selection factors, reflect slightly different filtering steps than for the datasets used in the DASM training (e.g., the validation data removes all sequences containing ambiguous nucleotides, while the training pipeline instead filters for conserved cysteines and removes parent–child pairs whose mutations occur only in ambiguous codons), as described in [17].

DASM is a transformer encoder that takes a parent amino acid sequence and outputs, at every site, a selection factor for each of the 20 amino acids [17]. The selection factor *f_j_*(*A → B*) is the multiplicative shift that selection applies to the probability of substituting *A* with *B* at site *j*, relative to the neutral expectation for that substitution. The DASM model we use to detect entrenchment, dasm 4m-v1jaffeCC+v1tangCC-joint, was trained as described in [17] on the Jaffe and Tang heavy-chain datasets; neither the training data nor the model input contains light chain or antigen sequence.

Thrifty supplies the neutral expectations against which those selection factors are defined. It is a convolutional network over learned 3-mer embeddings that predicts, for each nucleotide site and from local sequence context alone, a mutation rate and a conditional distribution over the three alternative bases [22]. Because it is trained on out-of-frame sequences, its predictions describe the mutation process rather than selection. The neutral model we use to validate DASM inferences with mutation rate ratios, ThriftyHumV0.2-59, was trained as described in [22].

All data required to reproduce the analyses in this paper, including the preprocessed repertoire datasets from [26, 17, 22], trained model weights, and pre-computed intermediate results, are available at Zenodo (https://doi.org/10.5281/zenodo.20171634). Analysis code and supplementary data files are available at https://github.com/matsengrp/dasm-epistasis-experiments, including a combined list of all identified entrenched amino acid substitutions and a summary table with DASM selection factors and rate-ratio validation results.

### Entrenchment detection using DASM selection factors

To identify entrenchment, we used DASM (Deep Amino acid Selection Model) [17] to predict selection factors on a dataset not used to train the model (Rodriguez). We used an unseen dataset to ensure that the selection factors represent generalizable predictions rather than overfitting artifacts that could produce spurious entrenchment signals. Because DASM predicts selection factors per sequence using context learned from its *∼*880,000-PCP training and validation set (Jaffe and Tang combined), the Rodriguez test set supports well-resolved per-site estimates despite being smaller than the training data. Given a parent sequence, DASM returns a selection factor for every amino acid at every site, so what we analyze is a prediction about that sequence rather than a tally of mutations observed in its descendants. We filtered sequences, sites, and selection factors so that:

- The parent sequence is the most recent common ancestor of each clonal family (i.e., root of the phylogenetic tree; parent–child pair depth = 2), yielding 12,942 PCPs for the Rodriguez dataset.
- Only sites where the parent amino acid is encoded by a germline codon are used.
- Target amino acid can be reached from parent sequence codon with a single-nucleotide substitution.

For each combination of site *j*, parent amino acid, target amino acid, and V family, we summarize selection by taking the median of the natural-log DASM selection factor *f* across sequences, requiring at least 10 observations per combination. For each site *j*, we considered reciprocal amino acid pairs (*A, B*) where both directions have an available median after the above steps:

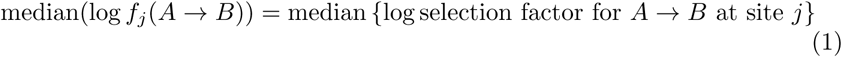

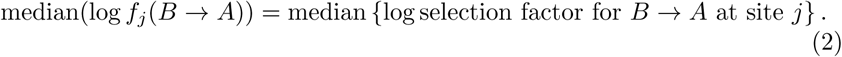

In figures, we denote median(log *f_j_*(*A → B*)) as “selection *A → B*.”

A site is classified as **entrenched** for the amino acid pair (*A, B*) if:

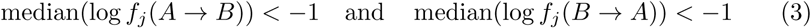

This criterion requires both reciprocal substitutions to occur at less than 37% of the neutral rate, indicating strong bidirectional purifying selection. Because we use only sites at which the parent amino acid is germline-encoded, the two medians come from disjoint sets of clonal families: those whose germline carries *A* at site *j*, and those whose germline carries *B*. A pair is therefore testable only if both amino acids occur in the germline at site *j*. Comparison can be made between different germline amino acids within a V family or between different V families.

### Validation of within-family grouping

Our entrenchment analysis aggregates selection factors across V gene alleles within a family that share the same germline amino acid at a given site. Because these V genes differ at up to *∼*20% of other positions, this pooling assumes that selection landscapes at a given site are similar across V genes in the group. To test this assumption, we computed median log selection factors for each V gene allele using the same filtering criteria as the family-level analysis. For each combination of site, parent amino acid, target amino acid, and individual V gene allele, we required *≥*3 observations. We assessed agreement using two metrics. First, the intraclass correlation coefficient (ICC), computed with a one-way random effects model partitioning variance into between-group (substitution identity) and within-group (V gene allele identity) components, was 0.97, 0.96, and 0.98 for IGHV1, IGHV3, and IGHV4 respectively. Second, threshold agreement (the fraction of individual V gene allele medians at family-level entrenched sites that independently fall below the *−*1 threshold) was 94%, 93%, and 92% for the three families. In the few cases of disagreement, dissenting V gene allele medians generally remained negative, indicating weaker purifying selection rather than a qualitative reversal, with the exceptions of site 9 in IGHV1 where two alleles of the same V gene (IGHV1-18) out of 21 showed positive medians and site 94 in IGHV3 where some V gene alleles showed medians near zero (Supplementary Figure S2). This validation applies equally to the between-family analysis, which relies on the same within-family aggregation.

### Solvent accessibility and backbone angle analysis

To characterize the structural context of entrenched sites, we analyzed antibody-antigen structures from the Structural Antibody Database (SAbDab) [19]. Structures were filtered to include only human antibodies with complete heavy chain, light chain, and antigen chains present, excluding heavy-chain-only antibodies (nanobodies, VHH). Structures with multiple antibodies were retained when V/J gene annotations were consistent across all entries for each PDB. We assigned V and J genes using ANARCI [35], and identified unmutated positions by comparison to germline references from the AIRR Community OGRDB (https://ogrdb.airr-community.org/germline_set/75).

We calculated solvent accessible surface area (SASA) using the DSSP algorithm [36] with Wilke reference values [37] to obtain relative solvent accessibility (RSA) on a 0–1 scale. To isolate the contributions of different binding partners to heavy-chain accessibility, we computed SASA in four scenarios: (1) the complete antibody-antigen complex (heavy + light + antigen chains), (2) the antibody alone (heavy + light chains), (3) the heavy chain in isolation, and (4) the heavy chain with the CDR-H3 loop removed. The antigen effect was calculated as the difference in RSA between the complete complex (scenario 1) and the antibody alone (scenario 2). The light-chain effect was calculated as the difference in RSA between the antibody alone (scenario 2) and the heavy chain in isolation (scenario 3). The CDR-H3 effect was calculated as the difference in RSA between the heavy chain in isolation (scenario 3) and the heavy chain with the CDR-H3 loop removed (scenario 4). We refer to these effects as ΔRSA; a positive value means the partner buries the residue.

We take the CDR-H3 to span positions 95–102, one position beyond the Chothia boundaries [18] at each end, so that the variable part of the V(D)J junction is included; the IGHV gene templates part of its first codon or two, though trimming during recombination varies from rearrangement to rearrangement [20, 21]. The adjacent FR4 (positions 103–113), though also encoded by the J gene segment, was retained in scenario 4 because the goal is to isolate the effect of the somatically diversified CDR-H3 loop; FR4 is a conserved framework segment, not part of this variable region.

We also counted atomic contacts in the same structures, as an alternative measure to ΔRSA, to confirm that the two agree. For each heavy-chain residue, we counted the non-hydrogen atoms of each partner (the antigen, the light chain, and the CDR-H3) within 5 Å of any non-hydrogen atom of the residue. For the CDR-H3, we excluded residues within two positions of the target residue in sequence, so that covalently bonded neighbors do not count as contacts. We compared the two measures in two ways. Visually, we compared their distributions across structures at each site. Quantitatively, for sites with at least 30 germline-encoded structures in a V family, we correlated the fraction of structures in which a partner lies within 5 Å with the fraction in which it buries the site (ΔRSA *>* 0). Both comparisons agree (Supplementary Figure S17).

Backbone dihedral angles (*ϕ* and *ψ*) were calculated using Biopython’s PP-Builder [38] to characterize local backbone geometry at each residue. For the dihedral analysis at site 9 we retained only structures carrying the germline amino acid at every position from 7 to 11, so that the conformations compared are not affected by somatic mutation within the window; this leaves 521 IGHV1 and 962 IGHV3 structures. All residue positions were numbered using the Chothia scheme [18].

### Grantham distance analysis

Grantham distance is a measure of physicochemical dissimilarity between amino acids based on composition, polarity, and molecular volume [39]. To test whether entrenched substitutions involve greater physicochemical change, we compared Grantham distances between entrenched and non-entrenched amino acid pairs. Because different sites have distinct amino acid repertoires, we used a stratified permutation test: within each V-family *×* site combination among the entrenched sites, we permuted the entrenched/non-entrenched labels (10,000 permutations), computed the per-stratum mean Grantham distance difference (entrenched minus non-entrenched), and averaged across strata. The one-sided *p*-value is the proportion of permuted statistics greater than or equal to the observed value. This asks whether entrenched pairs are physicochemically more distant than the other pairs available at the same site. It is not offered as evidence for entrenchment, but instead shows that entrenchment tracks physicochemical distance.

### Structural analysis of framework region 3 (FR3) hydrogen bonding networks

To investigate the structural basis of the observed entrenchment at site 73, we analyzed structures of Fab fragments from the Protein Data Bank (PDB). Structures were selected based on the following criteria: (1) human antibodies, (2) heavy-chain V gene assigned to either IGHV1 or IGHV3 family, (3) unmutated germline residues at positions 71–75, (4) presence of the entrenched amino acids at position 73, and (5) resolution *≤*3.0 Å. Criterion (3) ensures that the hydrogen bonding we report is not affected by somatic mutation within the loop. V gene assignments were performed using ANARCI [35], and unmutated positions were identified by comparison to germline references from the AIRR Community OGRDB (https://ogrdb.airr-community.org/germline_set/75). Four structures meeting these criteria were analyzed for each family: IGHV1 (PDB: 7X29, 2NY1, 6VY4, 8G3Z; resolution 1.99–2.49 Å) and IGHV3 (PDB: 4H8W, 3BN9, 6ULE, 6PPG; resolution 1.85–2.75 Å). Two of the IGHV1 structures, 7X29 and 8G3Z, were determined by cryo-electron microscopy and the remaining six by X-ray crystallography; the method is reported per structure in Supplementary Table S1. For each structure, PDB residue numbers were mapped to the Chothia numbering scheme by locating the conserved sequence motif at positions 71–75 (ADKST for IGHV1; RDNSK for IGHV3, or RDNAK where the germline V gene encodes alanine at position 74, as in 6PPG); mapping was verified individually per structure because PDB-to-Chothia offsets varied (direct correspondence in 6ULE, 4H8W, and 3BN9; +1 in 8G3Z, 7X29, 6VY4, and 6PPG; +3001 in 2NY1).

Structural analysis was performed using UCSF ChimeraX version 1.11.1 [40]. Because all heavy-chain copies within each analyzed structure have identical sequence, we analyzed a single heavy chain per structure to avoid redundancy. Hydrogen bonds involving sites 73–75 were identified using the hbonds command with two options: restrict any, which reports every H-bond in which the selected residues participate as either donor or acceptor against any partner (framework, CDRs, antigen), and intraRes false, which suppresses bonds within a single residue. Default geometric criteria (0.4 Å and 20*^◦^* tolerances relaxed from idealized per-atom-type geometry) give an effective absolute donor– acceptor distance cutoff of *∼*3.5 Å for N/O pairs. Each H-bond was classified by donor and acceptor type: backbone atoms are the amide N and carbonyl O, and all other donor/acceptor atoms are treated as sidechain.

### Density map figures

Density figures (Supplementary Figures S12B–C and S13) were prepared in ChimeraX as follows. Coordinates were fetched from the PDB and 2mFo–DFc maps from the PDBe electron density server (open <id= from eds); all maps were used as deposited. Displayed residues (sites 71–77 for Supplementary Figure S13; sites 7–11 for Supplementary Figure S12B–C) are shown in ball-and-stick, with cartoon retained nearby for context. In Supplementary Figure S13, hydrogen bonds are drawn using the hbonds criteria described above where their partners fall within the displayed region. Maps are displayed as transparent iso-surfaces contoured at 1.0*σ*, taken as the standard deviation of the full deposited map, and clipped to within 1.6 Å of the displayed residues (volume zone … range 1.6). Because contour appearance depends on resolution and map scaling, density is compared within panels rather than between them.

### CDR-H3 dependence of selection

We tested whether selection at a site depends on the CDR-H3 of the clonal family. To do so, we used the same filters as the entrenchment analysis: depth-2 pairs whose parent still carries the germline codon at the site, with the target amino acid one nucleotide away, restricted to IGHV1 and IGHV3 and to sites 1–94. We summarized each clonal family’s CDR-H3 by its length, because the structures make a concrete prediction about length: longer CDR-H3s bury entrenched sites more (Supplementary Figure S16C,D). We measured CDR-H3 length as the number of residues in that span.

For each substitution, we measured how much selection shifts between families with long and short CDR-H3s. A substitution is a site, its germline amino acid, and a target amino acid, within one V family. We split the clonal families of each V family at their median CDR-H3 length (13 residues for IGHV1, 12 for IGHV3; families at the median count as short), and took the difference in median log selection factor between the long and short halves, keeping substitutions with at least 50 clonal families across both halves. The difference of two sample medians is not zero even when the split carries no information, and the noise grows when families are few and their selection factors widely spread. We therefore estimated a noise floor for each substitution: we replaced CDR-H3 length with ten random family-level covariates, split at the median in the same way, and averaged the magnitude of the difference each produced. We subtracted this floor from the magnitude of the observed difference, kept the sign of the observed difference, and clipped at zero, so that a shift that does not exceed its floor counts as no shift.

We tested whether the shifts concentrate at the sites the CDR-H3 contacts. Contact frequency is the fraction of structures in which deleting the CDR-H3 increases that residue’s relative solvent accessibility, computed as in the solvent accessibility analysis above. We required at least 30 structures per site. We called a site a contact site if the CDR-H3 contacts it in more than half of that V family’s SAbDab structures. Substitutions at one site share clonal families and are not independent, so we tested at the site level, taking the median noise-corrected magnitude over the substitutions at each site. We tested magnitudes rather than signed differences, because a large shift in either direction means the CDR-H3 changed selection at a site; at the entrenched sites, where the structures predict the direction, we report the sign of each shift. We compared contact sites against all other sites with a one-sided Mann–Whitney test, analyzing IGHV1 and IGHV3 separately as independent replicates. Site 52 falls just below the contact cut in IGHV3 (49% of structures), so it joins the comparison group there, which works against the contrast.

For the structural comparison, we grouped structures by their own CDR-H3 length, keeping lengths carried by at least 20 structures, and averaged the burial of each entrenched site.

### Validation of DASM predictions using mutation rate ratios

To validate entrenchment predictions from DASM and its underlying neutral model Thrifty, we compared selection factors against mutation rate ratios calculated from observed versus expected mutation frequencies. Under the DASM framework, the selection factor *f_j_*(*A → B*) quantifies how selection shifts the probability of substitution *A → B* at site *j* relative to neutral expectation, given the parent amino acid sequence. When per-site mutation probabilities are small, substitution probabilities are approximately proportional to rates. The ratio of observed to expected mutation rates should therefore be approximately:

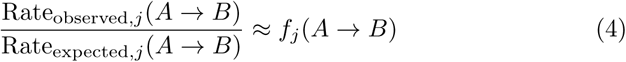

To estimate this ratio independently, we calculated Rate_observed_ from productive sequences, which have evolved under selection during affinity maturation, and Rate_expected_ from a baseline that is neutral with respect to amino acid changes [23, 22]. We used two complementary approaches for that baseline: mutation counts in out-of-frame sequences (Approach 1), which hypermutate without amino acid-level selection as described in “Data, code, and models” above, and mutation probabilities predicted by the Thrifty neutral model (Approach 2).

#### Mutation counts and rates

For both productive and out-of-frame datasets, we calculated mutation counts and rates from PCPs. We grouped all PCPs from the same V family, removing PCPs with ambiguous nucleotides (N) in their sequences. For productive sequences, which have undergone selection, we excluded leaf nodes to focus on mutations that have undergone subsequent rounds of selection [41]; for out-of-frame sequences, which evolve neutrally, we included all PCPs to maximize data. For each V family, site *j*, germline amino acid *A*, and target amino acid *B*, let Count(*A → B | j,* V family) denote the number of observed *A → B* substitutions across all PCPs where the parent sequence retains the germline amino acid *A* at site *j*.

To calculate rates from counts, we accounted for the different branch lengths of different PCPs (defined separately for each approach below):

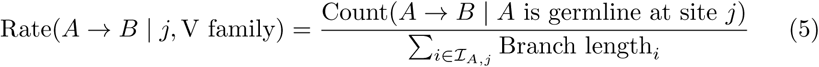

where *I_A,j_*denotes the set of all PCPs in the V family where the parent sequence has germline amino acid *A* at site *j*.

#### Approach 1: Out-of-frame data as neutral baseline

We calculated mutation rates separately for productive sequences (Rodriguez test dataset) and out-of-frame sequences (Tang-SHM dataset; Table 1). Although Tang-SHM is a held-out test dataset for Thrifty, Thrifty’s multi-hit correction component was trained using the Tang-SHM out-of-frame dataset, so Thrifty is not completely independent of the out-of-frame data. However, this multi-hit correction estimates only 3 parameters from the out-of-frame data, so the potential for data leakage affecting the validation is minimal. Out-of-frame sequences contain frameshifts in the V(D)J recombination region but remain in-frame within the V gene region, allowing Chothia numbering and germline annotation and thus ensuring compatibility across datasets.

To calculate productive and out-of-frame mutation rates, we calculated branch lengths for every PCP as the frequency of synonymous nucleotide mutations:

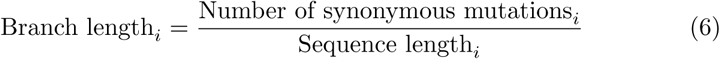

We chose synonymous mutation-based branch lengths because synonymous mutations are under more comparable selective pressure in both productive and out-of-frame datasets than nonsynonymous mutations, which experience differential selection. Although synonymous mutations are not a perfectly clean neutral proxy [22], any resulting bias approximately cancels because synonymous-based branch lengths are used symmetrically in the productive and out-of-frame rate calculations. We then calculated the mutation rate for each amino acid substitution using (5).

The rate ratio for each V family, site, and amino acid substitution is:

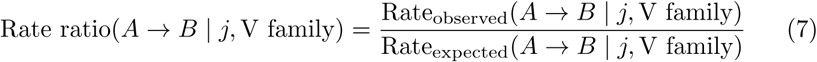

To stabilize estimates and avoid artifacts from sparse data, we applied pseudocount smoothing (adding 0.5 to mutation counts), and retained only substitutions with at least 5 mutations observed in the out-of-frame dataset. This filtering retained 1,180 out of 3,218 testable substitutions for validation.

#### Approach 2: Thrifty model as neutral baseline

As the size of the out-of-frame dataset limits validation coverage, we used a complementary approach that treats Thrifty’s neutral mutation model as ground truth to estimate expected counts. We used the much larger DASM model’s training dataset (Jaffe and Tang, Table 1) for this approach to increase statistical power. This dataset was used to train DASM, so this validation tests the DASM model but does not test whether its predictions generalize to unseen data.

To calculate Thrifty-predicted expected counts, we:

1. Calculated branch lengths as the total nucleotide mutation frequency multiplied by a scaling factor *α* = 1.6:

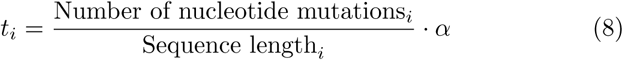

where *α* is derived from regressing observed mutation frequencies against DASM-optimized branch lengths during DASM training. This correction accounts for purifying selection, which removes some expected mutations (1*/*1.6 *≈* 0.63, meaning observed mutations are approximately 63% as likely as neutral expectation). Because *α* is a single global scalar, it adjusts the overall scale of expected counts but cannot introduce a correlation where there is not one.
2. Converted Thrifty’s neutral mutation rates 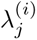^)^ at each site *j* for each PCP *i* into mutation probabilities: 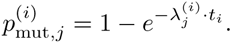
3. Multiplied by Thrifty’s conditional substitution probabilities *P*_Thrifty_(*B | A*; *j, i*) and summed across the same PCPs used to count observed mutations:

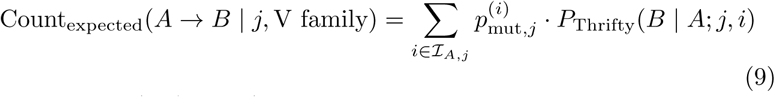

where *P*_Thrifty_(*B | A*; *j, i*) is obtained by summing Thrifty’s nucleotide-level conditional substitution probabilities over codons encoding amino acid *B*, and *I_A,j_*is defined as in (5).

Since both observed and Thrifty-predicted expected counts are calculated on the same phylogenetic trees and per-site mutation probabilities are small, branch lengths approximately cancel when taking ratios, allowing direct comparison of mutation counts rather than rates:

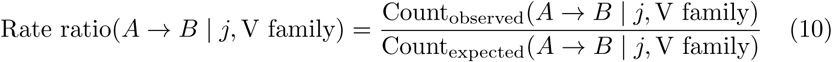

We applied the same pseudocount smoothing (0.5) to both observed and expected counts and the same filtering criteria (expected mutations *≥* 5), then compared observed productive mutation counts to Thrifty-predicted expected counts. This approach retained 2,877 out of 3,247 testable substitutions for validation.

#### Comparison to DASM selection factors

For both validation approaches, we compared log-transformed rate/count ratios to median log DASM selection factors, aggregated using the same filtering and summarization criteria described in “Entrenchment detection using DASM selection factors” above (depth 2 PCPs, germline-codon parents, single-nucleotide-reachable targets, *≥*10 observations, median per combination). We assessed agreement using Pearson correlation coefficients and orthogonal regression (Deming, *λ* = 1), which is more appropriate than ordinary least squares regression when both variables have measurement error. To assess whether entrenched sites showed consistent purifying selection, we verified that substitutions identified as entrenched fell in the purifying selection region (log ratio *<* 0) of the validation plots.

### Germline allele misassignment analysis

Closely related V alleles can be confused during germline annotation when somatic hypermutation hits a site where they differ, so an amino acid acquired by somatic mutation can be mistaken for a germline residue. Such misassignment does not alter the selection factors themselves, which DASM computes from the root sequence alone without annotation, but it can place the sequence in the wrong germline amino acid group in the entrenchment analysis. Misassignment thus contaminates a germline group with sequences that acquired the amino acid by somatic mutation before the clonal family root. Entrenchment in a contaminated group may therefore reflect something other than selection on the germline residue. To test how often misassignment happens at entrenched sites, we simulated clonal family root sequences (most recent common ancestors) with known V alleles and measured how often germline annotation assigns an allele that encodes a different amino acid there. For each naive-to-root parent–child pair in the Rodriguez dataset whose clonal family contributes to the entrenchment analysis, we simulated 10 replicate root sequences, starting from the naive sequence and evolving it along the branch length, with Thrifty for neutral mutation and DASM for selection. The naive sequences are inferred rather than observed, but they are the known starting point of the simulation and so serve as ground truth. The simulated root sequences closely match the real root sequences in both total mutation count and per-codon mutation frequency along the V gene (Supplementary Figure S15A–B). We then annotated all simulated sequences with partis [20], the tool we used to annotate the real data, and compared the assigned V allele to the true (simulation-input) allele. The comparison is not circular, since partis uses neither DASM nor Thrifty. Overall, 99.5% of simulated sequences were assigned the correct V gene and 92.4% the correct V allele. For each V family, site, and assigned germline amino acid, the contamination rate is the fraction of simulated sequences whose assigned allele encodes a different amino acid at that site than the true allele. We annotated the simulated sequences against the full default germline set rather than per-subject inferred genotypes, so partis chose among more candidate alleles than in the annotation of the real data, and these rates likely overestimate misassignment.

Three entrenched site/amino acid combinations have substantial contamination (Supplementary Figure S15C), all traceable to specific allele confusions: IGHV1 site 50 R (22.5%, mostly from IGHV1-69 and partly from IGHV1-2 allele pairs), IGHV1 site 73 K (36.6%, from IGHV1-69 allele pairs), and IGHV4 site 29 V (10.6%, from IGHV4-59/61 confusion). The other entrenched amino acids at site 50 in both IGHV1 and IGHV3 have negligible contamination, so the entrenchment signal at this site does not depend on the contaminated group. At site 73, most contamination comes from sequences whose true allele is IGHV169*01 (E) but which are assigned IGHV1-69*06 (K), an allele that differs from *01 only at this site. The other K-carrying alleles in our data (IGHV1-69*02, *04, and *10) are each at least three nucleotides from any E-carrying allele, and so are much less prone to this confusion; each has a median log selection factor below *−*1 for K*→*N (Supplementary Figure S2), so entrenchment at this site holds for germline-encoded K, not only for K acquired by somatic mutation. Site 29 V is observed in only one allele in our data (IGHV4-61*01), so there we cannot distinguish germline-encoded entrenchment from entrenchment of somatic mutations acquired before the clonal family root. All remaining entrenched site/amino acid combinations have contamination below 4%, with most below 1%, indicating that misassignment does not drive entrenchment at the large majority of sites.

## Acknowledgements

We thank Phil Bradley for helpful discussions.

This work was supported by NIH grant R01-AI146028 (PI Matsen). Scientific Computing Infrastructure at Fred Hutch is funded by ORIP grant S10OD028685. Frederick Matsen is an investigator of the Howard Hughes Medical Institute.

## Data Availability Statement

The code for reproducing this analysis and the supplementary data files are available at https://github.com/matsengrp/dasm-epistasis-experiments. The preprocessed data and the trained DASM model used in this paper are available in Zenodo at https://doi.org/10.5281/zenodo.20171634.

## Supplementary Materials

**Figure S1:**
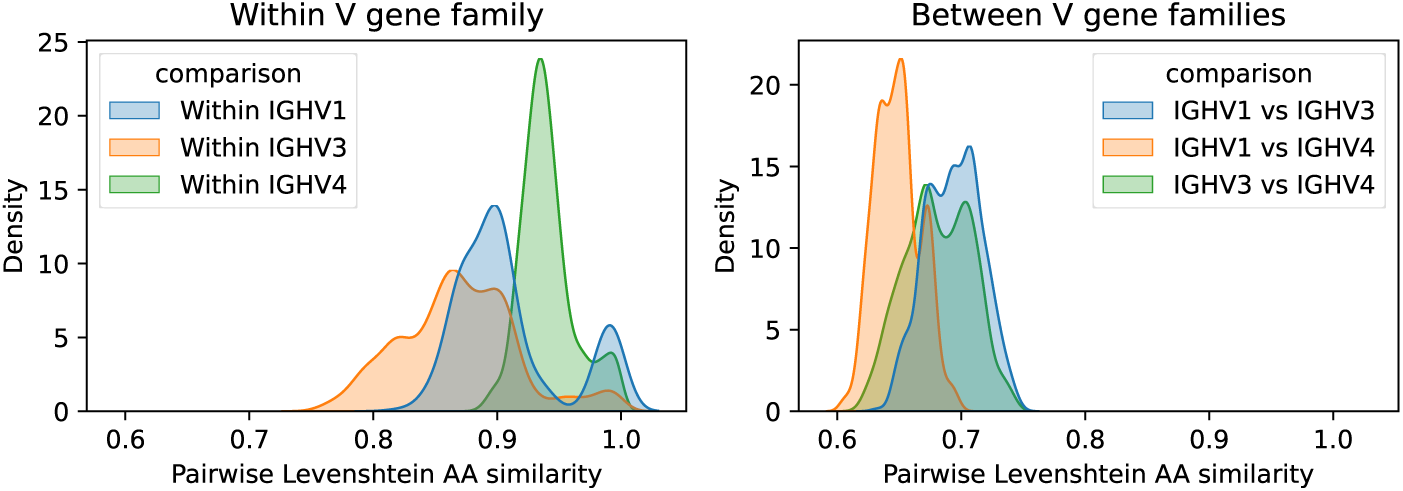
Pairwise amino acid similarity between IGHV genes. Kernel density estimates of pairwise Levenshtein amino acid similarity for IGHV1, IGHV3, and IGHV4. Top: within-family comparisons, where most gene pairs differ at up to *∼*10% (IGHV4), *∼*15% (IGHV1), or *∼*20% (IGHV3). Bottom: between-family comparisons, where pairs differ at 25–40%.

**Figure S2:**
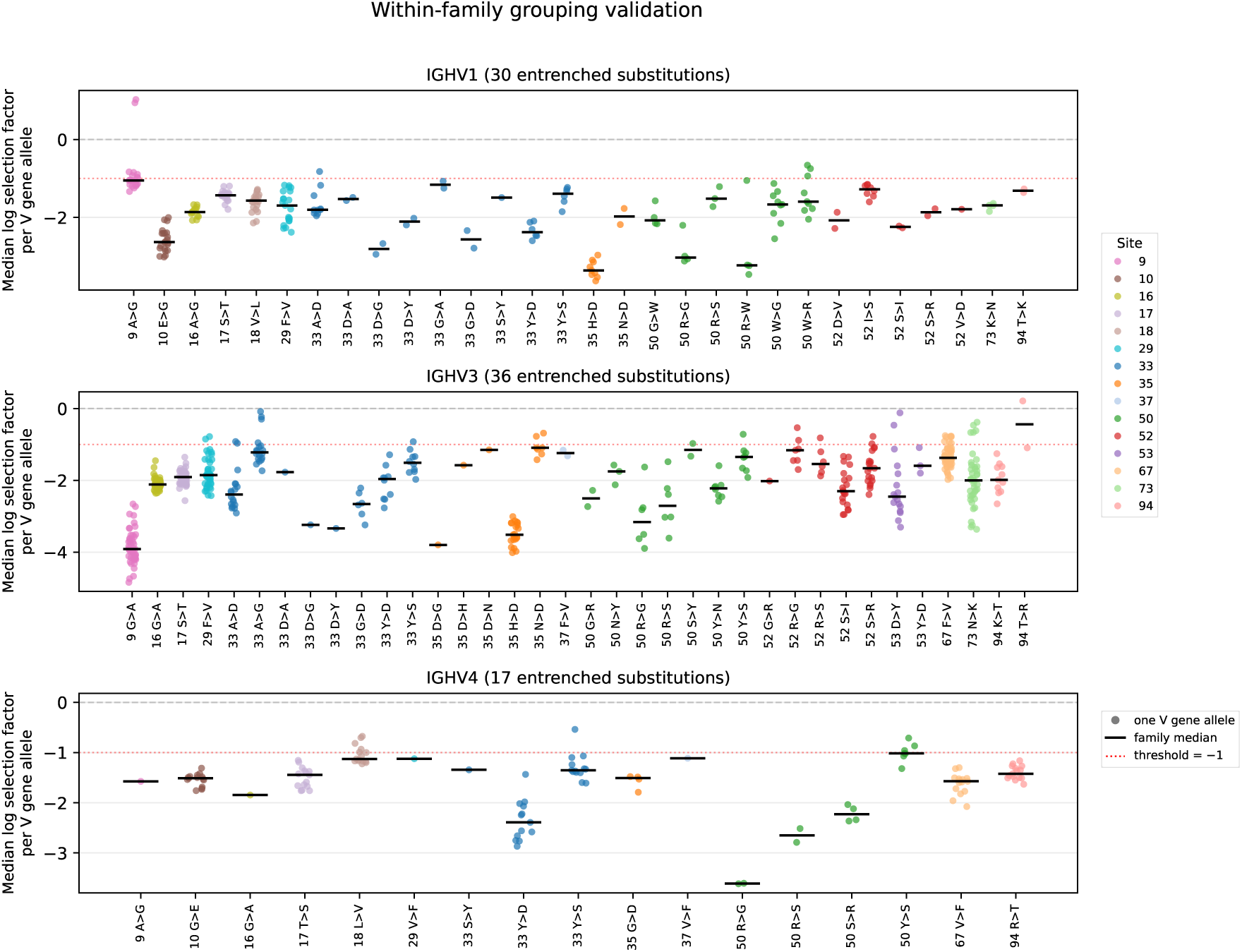
Within-family grouping validation: per-V-gene-allele median selection factors at entrenched sites from both within-family and between-family analyses. Each dot represents the median log selection factor for one V gene allele at one entrenched substitution, colored by site. Black bars show the family-level median used in the main analysis. The dashed red line indicates the *−*1 entrenchment threshold. Per-V-gene-allele medians cluster tightly around the family median, with the exceptions noted in Methods (site 9 in IGHV1, site 94 in IGHV3).

**Figure S3:**
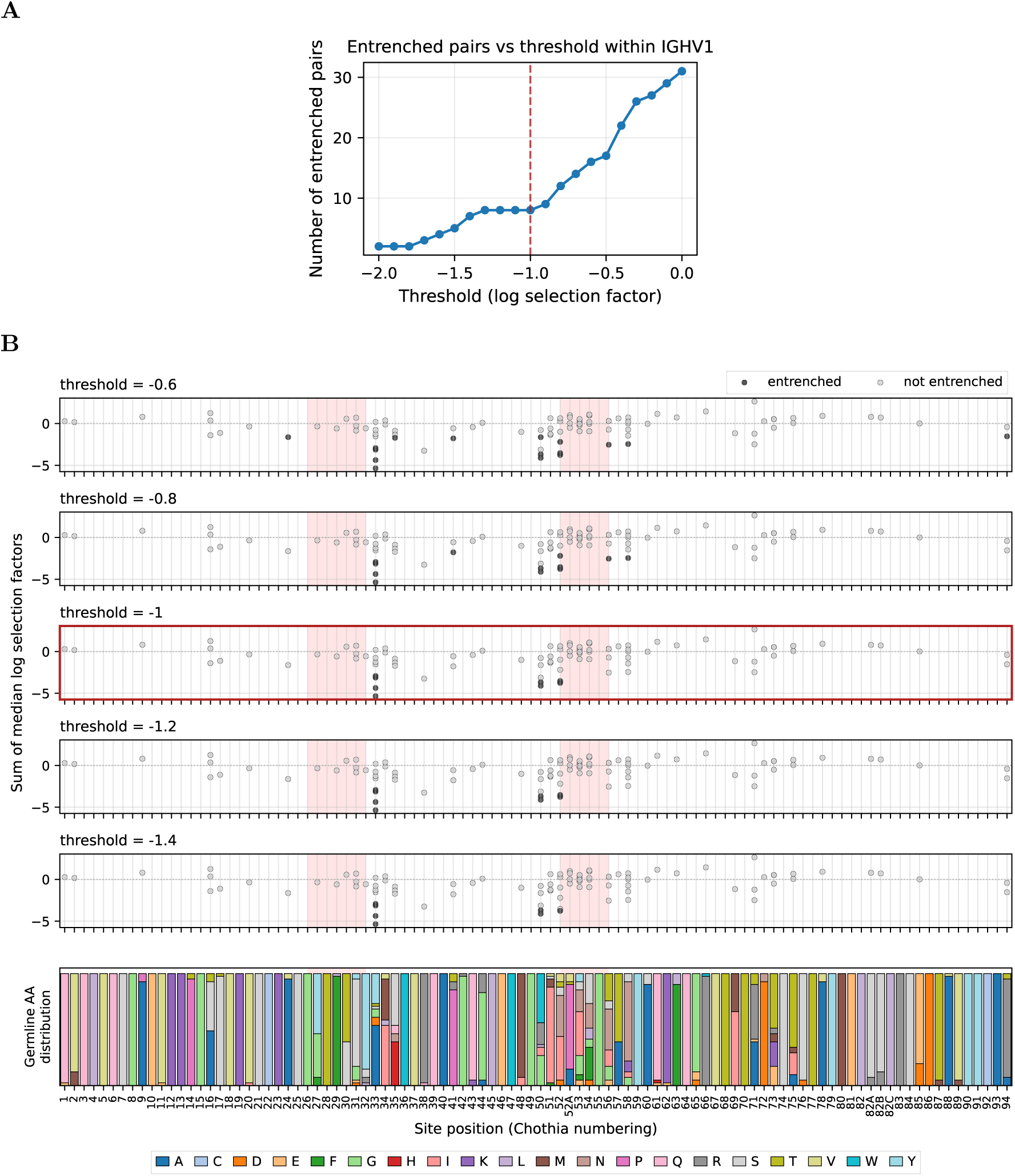
Threshold sensitivity of entrenchment calls within IGHV1. **(A)** Number of entrenched amino acid pairs as a function of the log selection factor threshold. The dashed red line marks the chosen cutoff of *−*1, at which we identify 8 entrenched pairs at 3 sites. Tightening to *−*1.4 retains 7 of these 8 pairs and all 3 sites. **(B)** Per-site view at five thresholds (top to bottom: *−*0.6, *−*0.8, *−*1, *−*1.2, *−*1.4). Filled points mark entrenched amino acid pairs, open points mark non-entrenched pairs; pink shading highlights CDR regions. The bottom track shows the germline amino acid distribution across V genes at each site.

**Figure S4:**
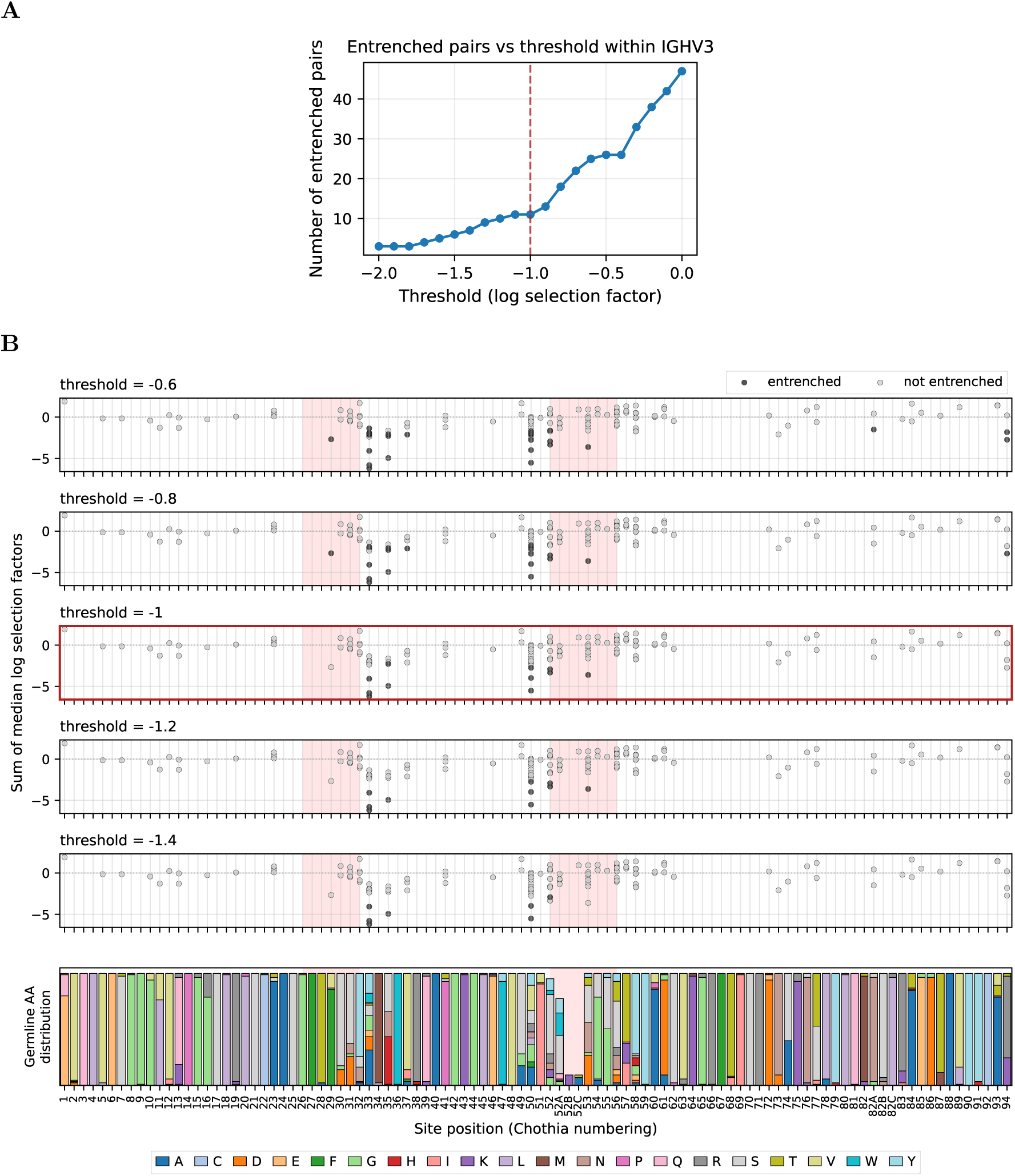
Threshold sensitivity of entrenchment calls within IGHV3. (A) Number of entrenched amino acid pairs as a function of the log selection factor threshold. The dashed red line marks the chosen cutoff of *−*1, at which we identify 11 entrenched pairs at 5 sites. Tightening to *−*1.4 retains 7 of these 11 pairs and 4 of the 5 sites. **(B)** As in Supplementary Figure S3B, per-site view at five thresholds (top to bottom: *−*0.6, *−*0.8, *−*1, *−*1.2, *−*1.4).

**Figure S5:**
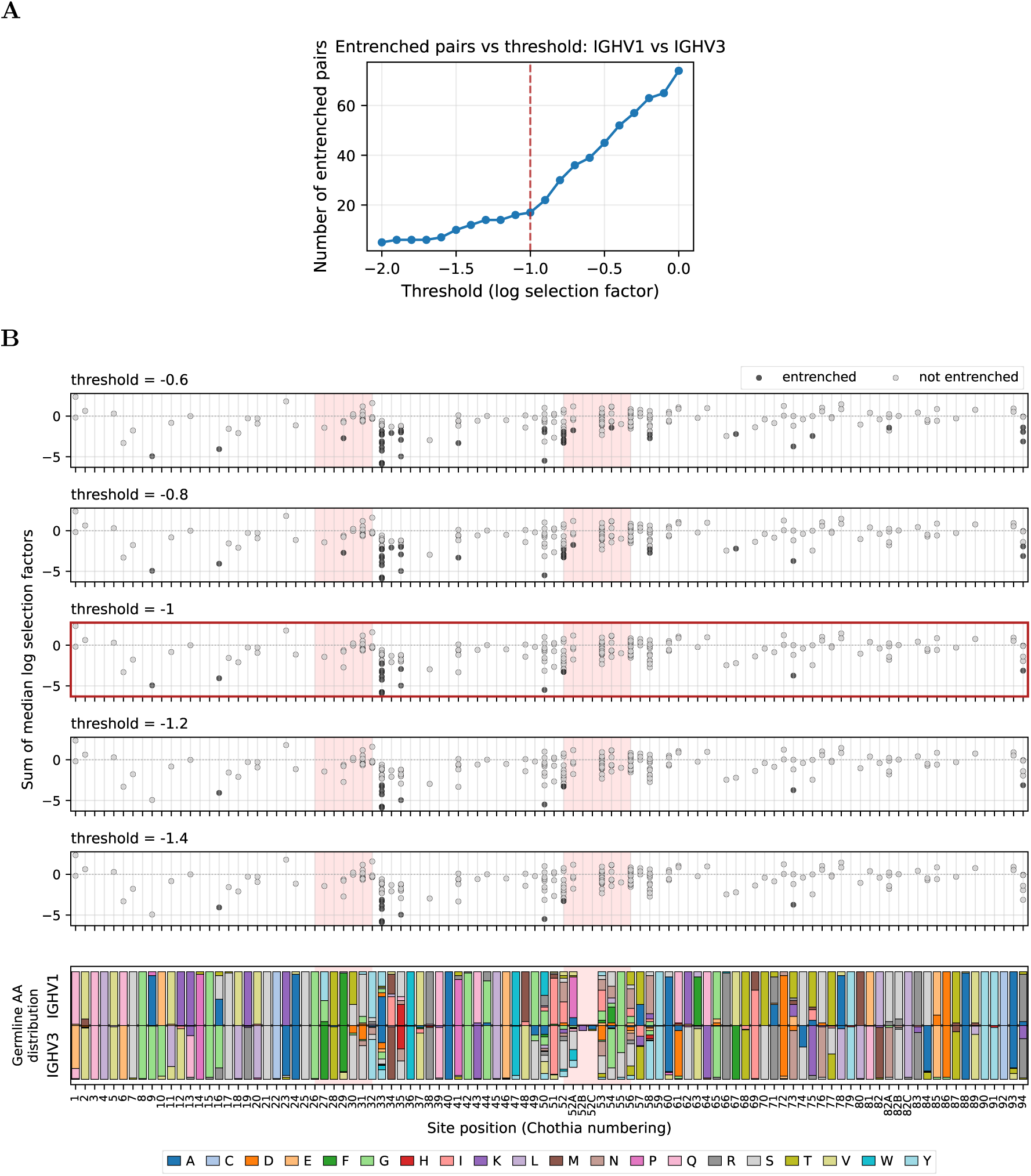
Threshold sensitivity of entrenchment calls between IGHV1 and IGHV3. **(A)** Number of entrenched amino acid pairs as a function of the log selection factor threshold. The dashed red line marks the chosen cutoff of *−*1, at which we identify 17 entrenched pairs at 8 sites. Tightening to *−*1.4 retains 12 of these 17 pairs and 6 of the 8 sites. **(B)** As in Supplementary Figure S3B, per-site view at five thresholds (top to bottom: *−*0.6, *−*0.8, *−*1, *−*1.2, *−*1.4).

**Figure S6:**
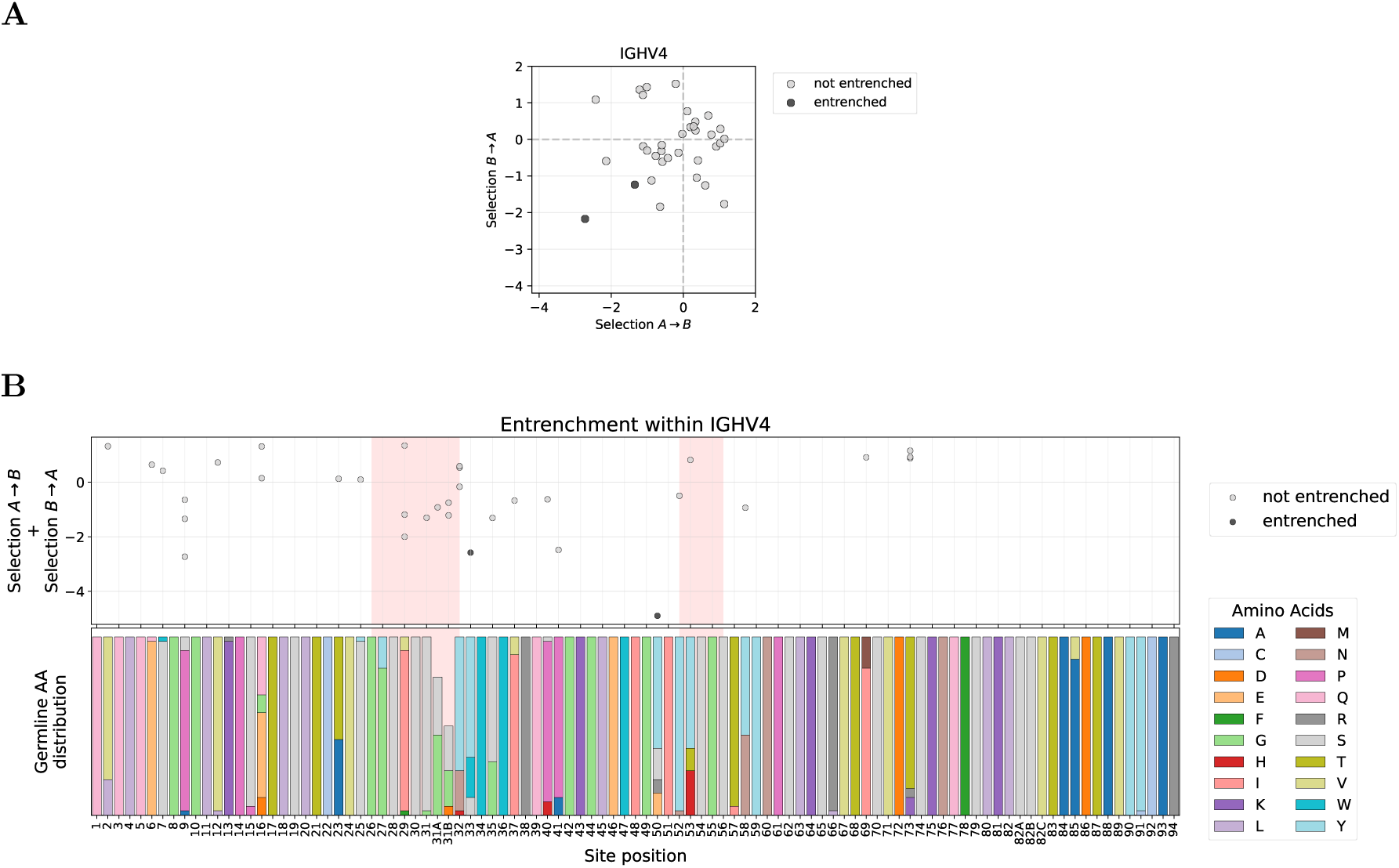
Within-family entrenchment analysis for IGHV4. **(A)** Reciprocal median log selection factors (selection *A → B* and selection *B → A*) for amino acid pairs at sites where V genes within IGHV4 differ in germline identity; entrenched pairs (both reciprocal median log selection factors *< −*1) are highlighted. **(B)** Entrenchment mapped along the V gene sequence, analogous to Figure 2B. IGHV4 shows recurrent entrenched sites consistent with those identified in IGHV1 and IGHV3.

**Figure S7.**
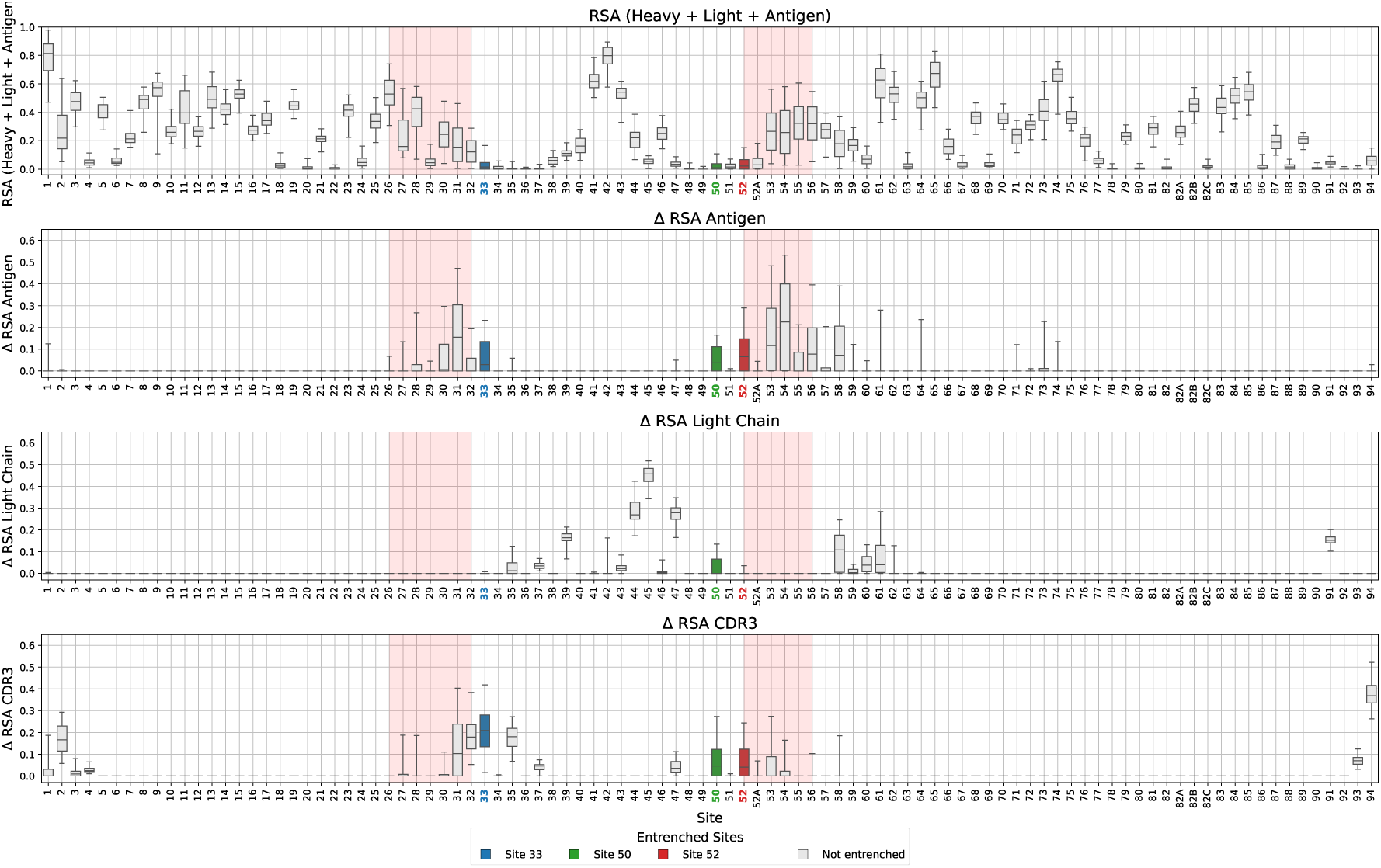
RSA analysis for IGHV1 family, with within-family entrenched sites high-lighted. Row 1: Relative solvent accessibility (RSA) in the full complex. Row 2: Change in RSA upon antigen removal (antigen effect). Row 3: Change in RSA upon light-chain removal (light-chain effect). Row 4: Change in RSA upon CDR-H3 removal. This is the IGHV1 counterpart to the IGHV3 analysis shown in Figure 3D.

**Figure S8:**
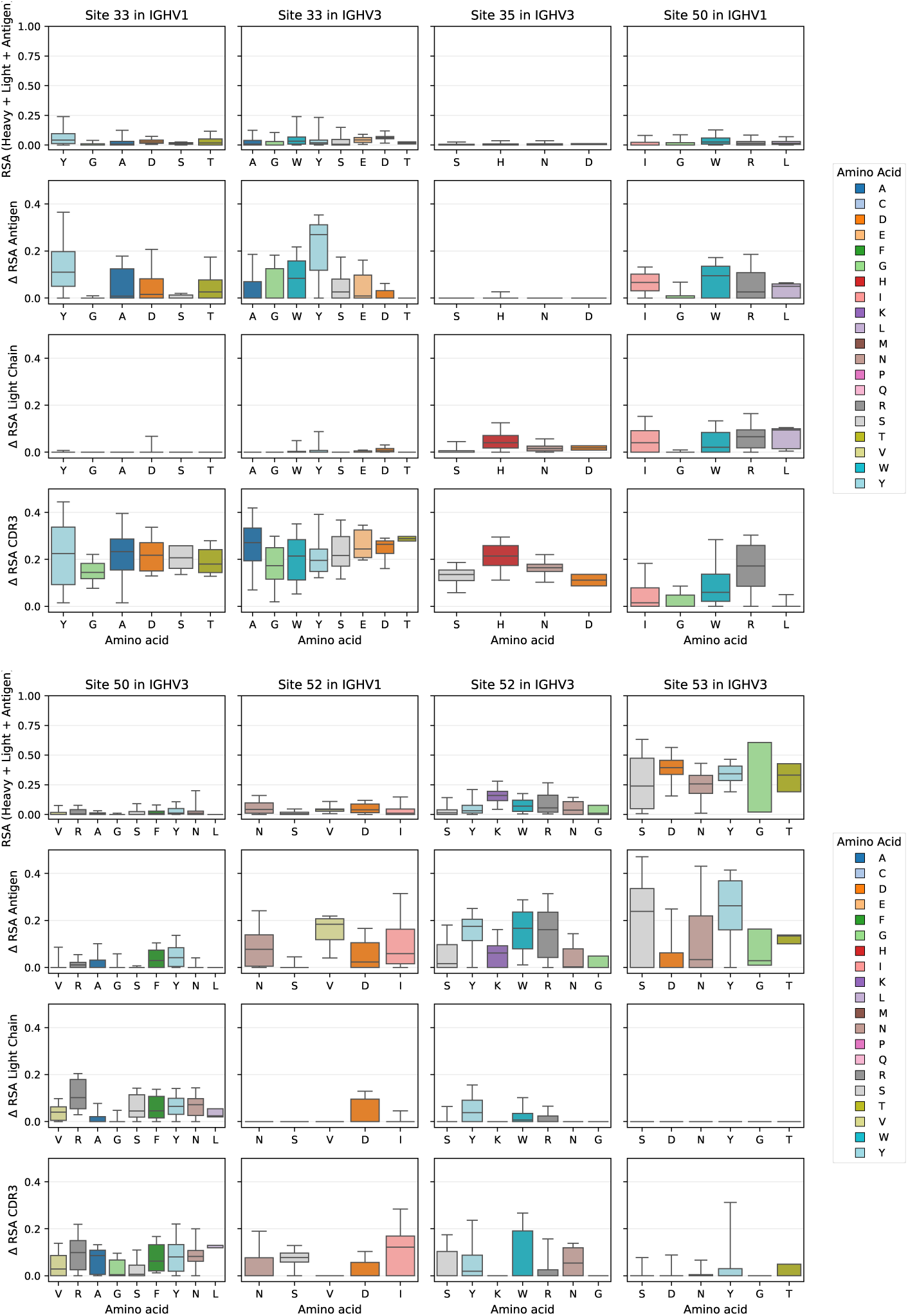
RSA properties at entrenched sites by germline amino acid. RSA properties for all within-family entrenched sites in IGHV1 and IGHV3. Row one shows RSA of the amino acid in the full heavy chain, light chain, and antigen complex. Row two shows the change in RSA upon antigen removal (antigen burial). Row three shows the change in RSA upon light-chain removal (light-chain burial). Row four shows the change in RSA upon CDR-H3 removal.

**Figure S9:**
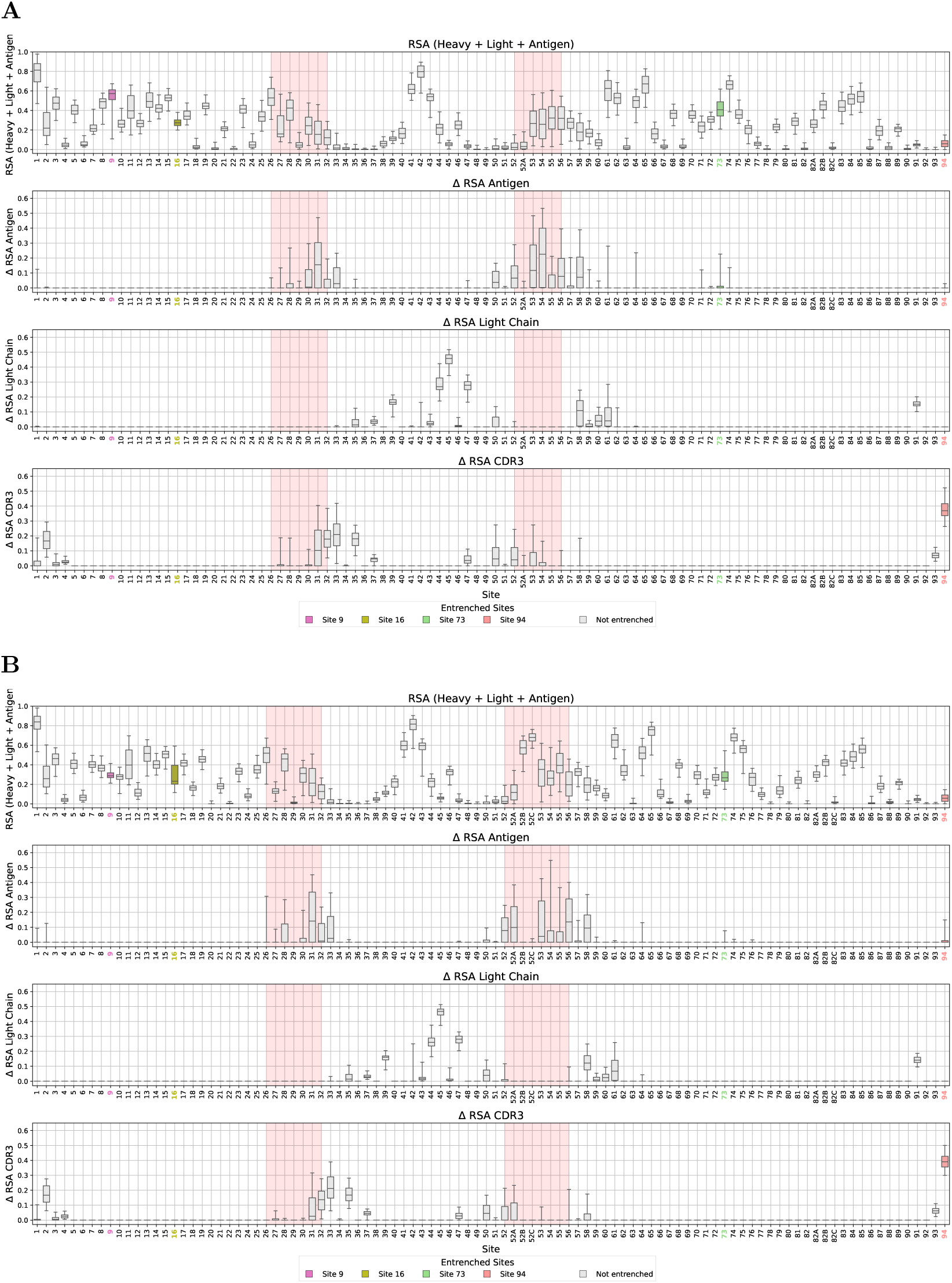
RSA analysis with sites entrenched between V families but not within V families (newly added sites) highlighted. **(A)** IGHV1 family: relative solvent accessibility (RSA) of sites in the full antibody-antigen complex, and change in RSA upon removal of antigen, light chain, or CDR-H3. **(B)** IGHV3 family: same analysis as (A).

**Figure S10:**
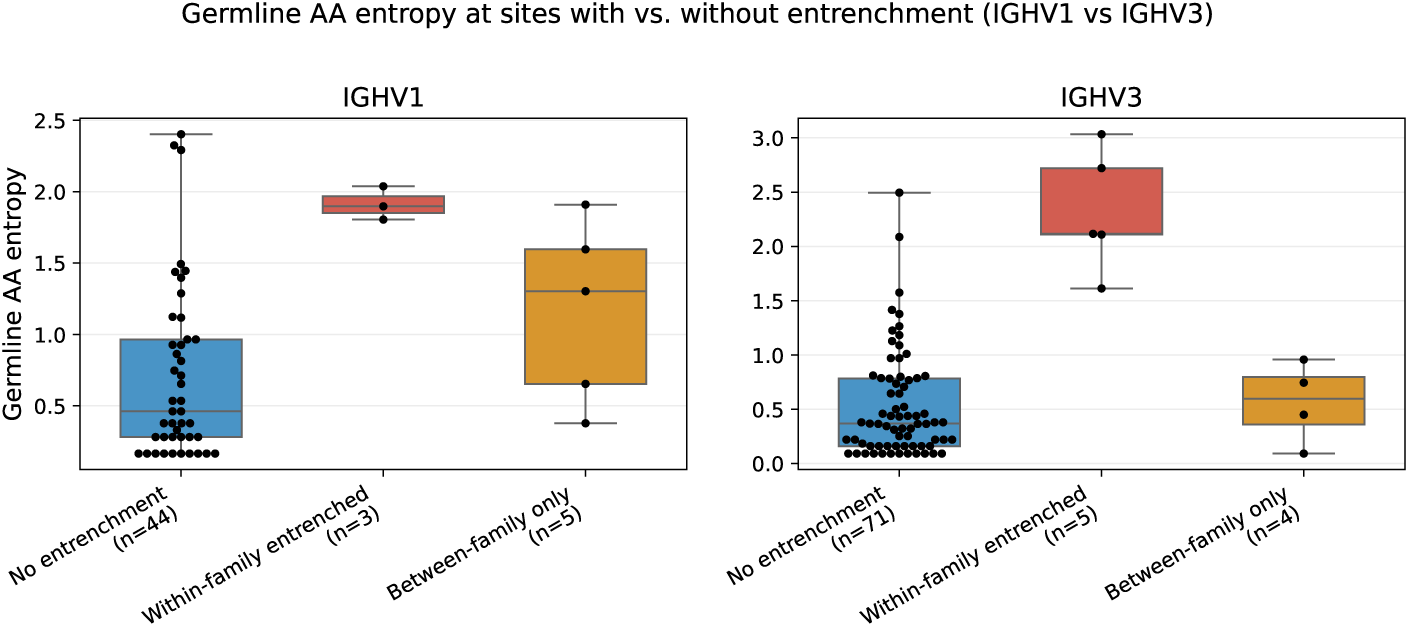
Shannon entropy for sites entrenched within and between IGHV1 and IGHV3. Germline diversity (Shannon entropy) at sites classified as non-entrenched, within-family entrenched, or between-family entrenched (excluding sites that overlap with within-family entrenched). Between-family-only entrenched sites show lower germline diversity than within-family entrenched sites, consistent with structural rather than diversifying constraints. Sites with only one germline amino acid across V genes in the family (entropy = 0) are excluded.

**Figure S11:**
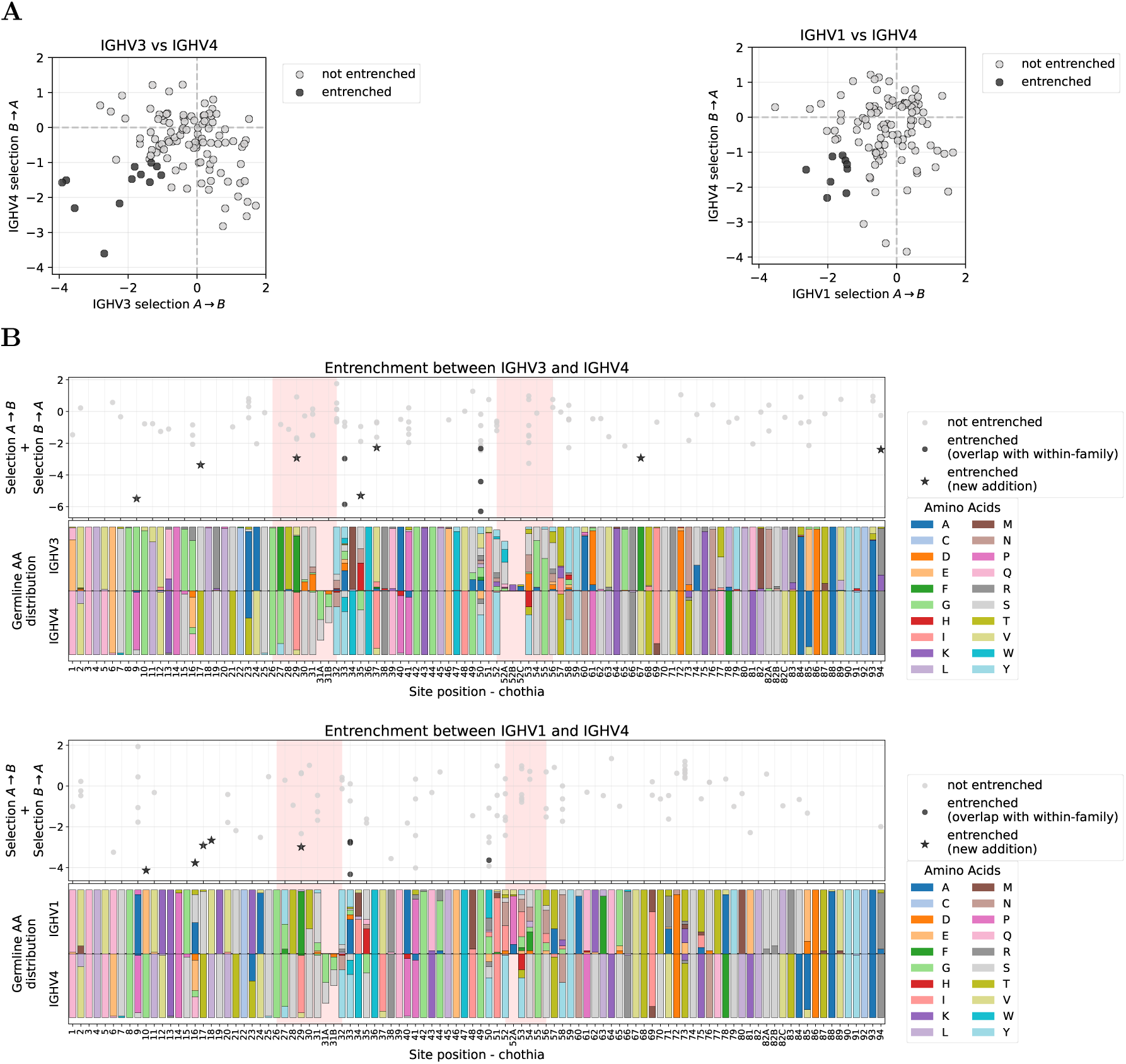
Between-family entrenchment analysis for additional V-family pairs. Panels show results for IGHV3 vs IGHV4 (left in A, top in B) and IGHV1 vs IGHV4 (right in A, bottom in B). **(A)** Reciprocal median log selection factors (selection *A → B* vs. selection *B → A*) for amino acid pairs at sites where the two families differ in germline identity; entrenched pairs (both *< −*1) are highlighted. **(B)** Entrenchment mapped along the V gene sequence with within-family overlay, analogous to Figure 5B.

**Figure S12:**
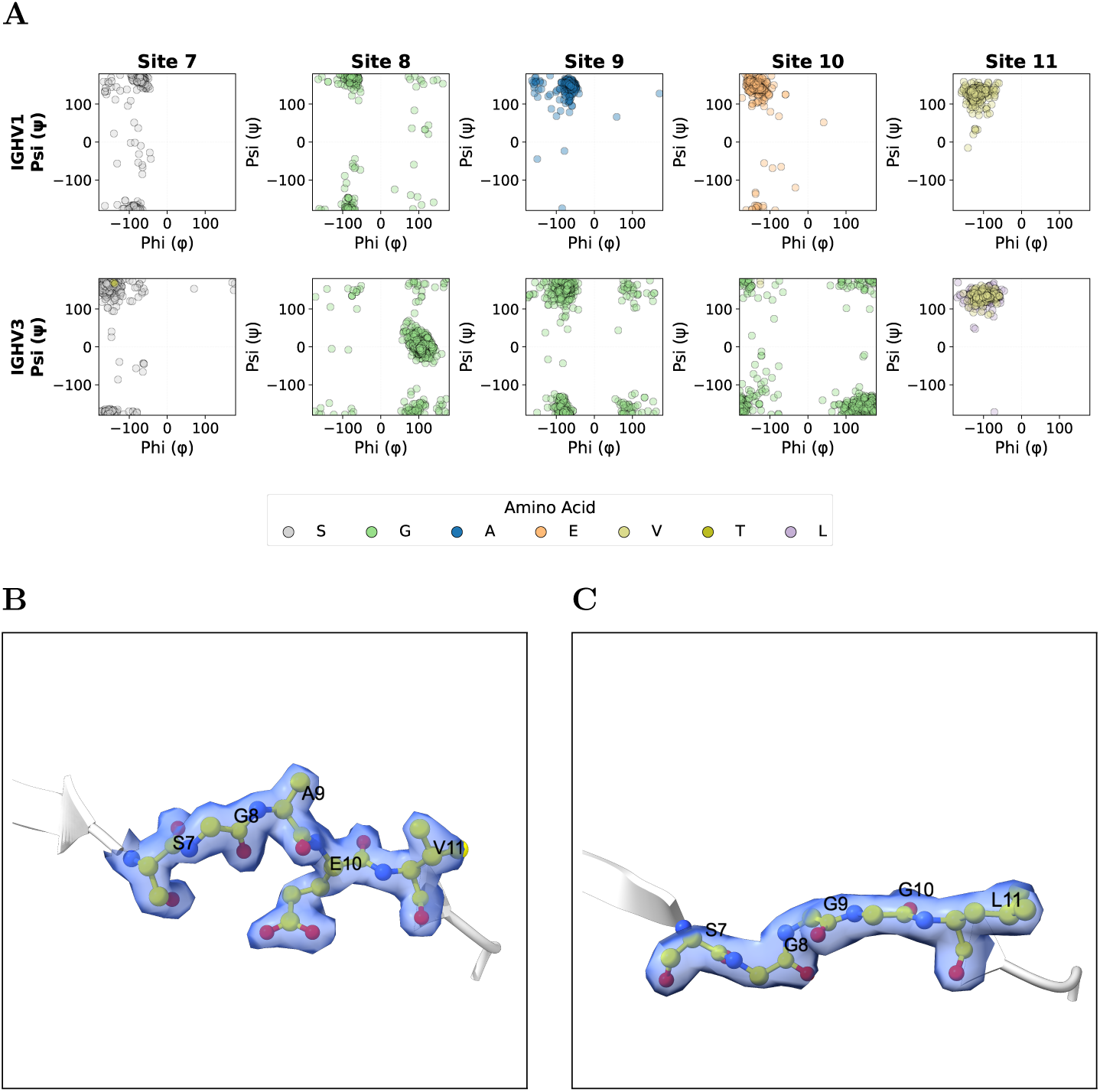
Backbone data across SAbDab for entrenchment at site 9. **(A)** Ramachandran plots for sites 7–11 showing backbone dihedral angles (phi and psi) in IGHV1 (top) and IGHV3 (bottom) across SAbDab structures. Structures are included only if they carry the germline amino acid at every site in the 7–11 window, giving 521 IGHV1 and 962 IGHV3 structures. The distinct angle distributions confirm family-specific backbone conformations at the site 9 region. **(B, C)** Density at sites 7–11 for the two structures shown in Figure 6A: **(B)** IGHV1, 7VYT (chain H, X-ray, 1.53 Å) and **(C)** IGHV3, 8FFE (chain H, X-ray, 1.72 Å), both 2mFo–DFc at 1.0*σ*. Density is shown as a transparent surface around the backbone. In both structures the 7–11 backbone is continuously enclosed by density, so the differing conformations in Figure 6A are supported by the maps rather than by modeling alone.

**Table S1:**
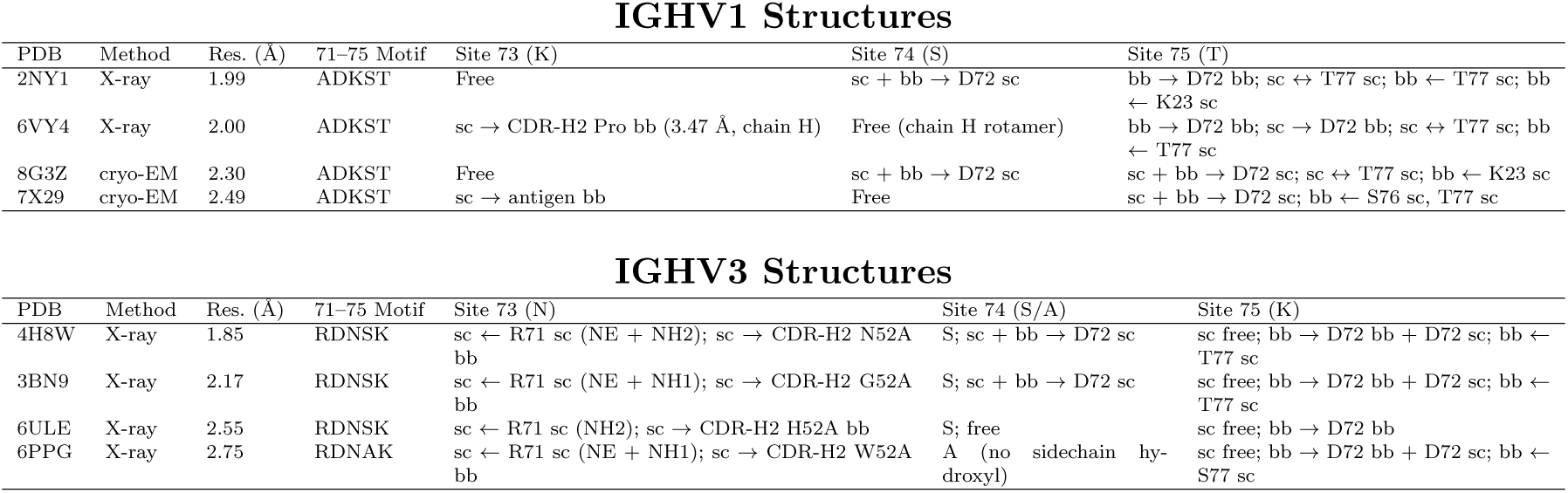
Structural analysis of hydrogen bonding networks at loop 73–75 in IGHV1 and IGHV3 Fab structures. Each row summarizes the local H-bond interactions observed at the loop containing entrenched site 73 for one structure, derived from ChimeraX hbonds analysis with default geometric criteria (effective *∼*3.5 Å donor–acceptor cutoff for N/O pairs). All structures carry the unmutated germline motif at sites 71–75, so the interactions reported here are not the result of somatic mutation within the loop. In IGHV1, K73 has no consistent framework hydrogen bond partner, while T75 is anchored by hydrogen bonds to both D72 and T77 in all four structures. In IGHV3, N73 is anchored by two contacts present in all four structures: a hydrogen bond from the R71 guanidinium and a donor hydrogen bond to a CDR-H2 backbone carbonyl (the acceptor lies at position 52A in all four structures, though its identity varies: His, Asn, Gly, or Trp); the K75 sidechain is free of hydrogen bonds. Notation key: each H-bond is annotated as sidechain (sc) or backbone (bb) on each side (backbone = amide N and carbonyl O). Specific atom names appear in parentheses only to disambiguate multiple contacts on the same arginine: NE is the *ε* guanidinium nitrogen and NH1/NH2 are the two terminal guanidinium nitrogens. All are sidechain atoms.

| IGHV1 Structures |  |  |  |  |  |  |
|---|---|---|---|---|---|---|
| PDB | Method | Res. (Å) | 71–75 Motif | Site 73 (K) | Site 74 (S) | Site 75 (T) |
| 2NY1 | X-ray | 1.99 | ADKST | Free | sc + bb → D72 sc | bb → D72 bb; sc ↔ T77 sc; bb ← T77 sc; bb ← K23 sc |
| 6VY4 | X-ray | 2.00 | ADKST | sc → CDR-H2 Pro bb (3.47 Å, chain H) | Free (chain H rotamer) | bb → D72 bb; sc → D72 bb; sc ↔ T77 sc; bb ← T77 sc |
| 8G3Z | cryo-EM | 2.30 | ADKST | Free | sc + bb → D72 sc | sc + bb → D72 sc; sc ↔ T77 sc; bb ← K23 sc |
| 7X29 | cryo-EM | 2.49 | ADKST | sc → antigen bb | Free | sc + bb → D72 sc; bb ← S76 sc, T77 sc |
| IGHV3 Structures |  |  |  |  |  |  |
| PDB | Method | Res. (Å) | 71–75 Motif | Site 73 (N) | Site 74 (S/A) | Site 75 (K) |
| 4H8W | X-ray | 1.85 | RDNSK | sc ← R71 sc (NE + NH2); sc → CDR-H2 N52A bb | S; sc + bb → D72 sc | sc free; bb → D72 bb + D72 sc; bb ← T77 sc |
| 3BN9 | X-ray | 2.17 | RDNSK | sc ← R71 sc (NE + NH1); sc → CDR-H2 G52A bb | S; sc + bb → D72 sc | sc free; bb → D72 bb + D72 sc; bb ← T77 sc |
| 6ULE | X-ray | 2.55 | RDNSK | sc ← R71 sc (NH2); sc → CDR-H2 H52A bb | S; free | sc free; bb → D72 bb |
| 6PPG | X-ray | 2.75 | RDNAK | sc ← R71 sc (NE + NH1); sc → CDR-H2 W52A bb | A (no sidechain hy-droxyl) | sc free; bb → D72 bb + D72 sc; bb ← S77 sc |

**Figure S13:**
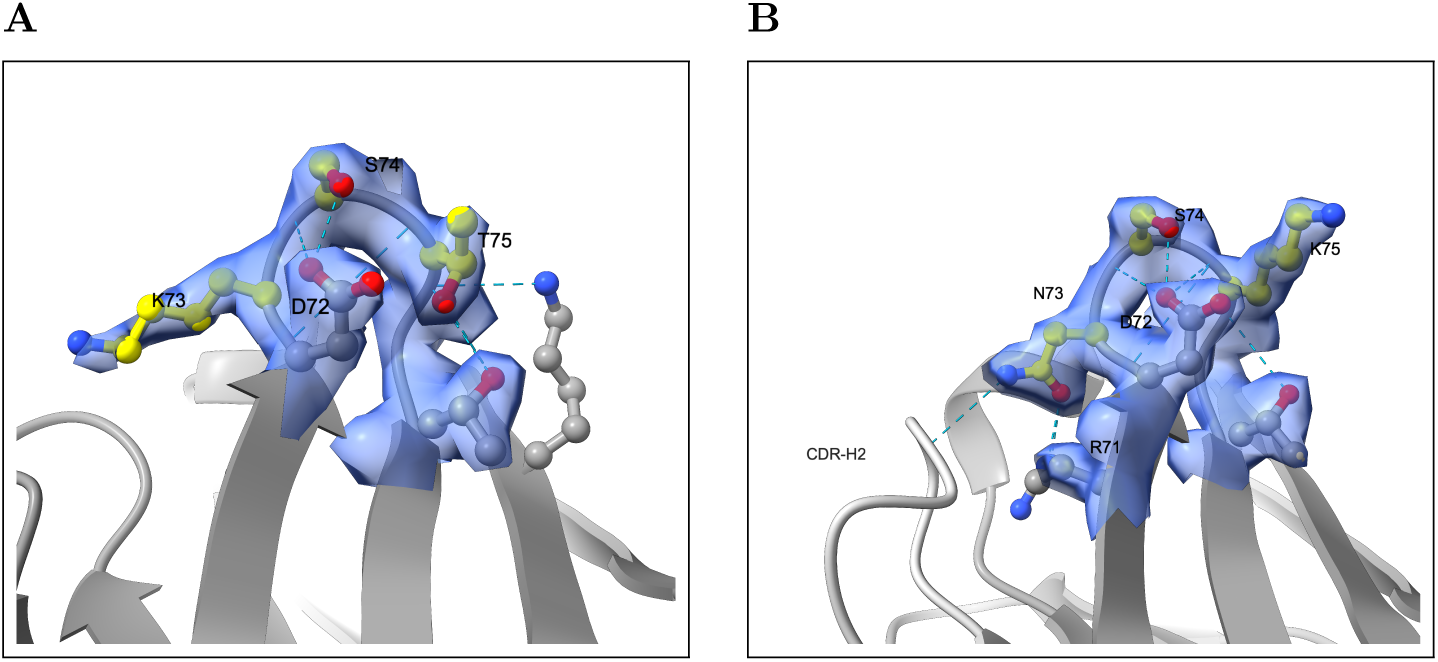
Density support for Figure 6B–C: site 73 in IGHV1 and IGHV3. Both panels show the same structures as Figure 6B–C, with 2mFo– DFc density drawn as a transparent surface around the loop; sites 73–75 are yellow, hydrogen bond partners grey, and hydrogen bonds cyan dashes. Supplementary Table S1 reports the hydrogen bond analysis for these structures. **(A)** IGHV1, 2NY1 (chain D, X-ray, 1.99 Å), contoured at 1.0*σ*. All nine atoms of the K73 residue are modeled. Density encloses the backbone and proximal sidechain through C*δ*, but falls below the 1.0*σ* contour at the terminal two atoms, C*ε* and N*ζ*. The sidechain extends into solvent rather than toward the framework, and its terminus is correspondingly less well enclosed than the framework residues around it, as expected of a sidechain with no fixed hydrogen bond partner. **(B)** IGHV3, 4H8W (chain H, X-ray, 1.85 Å), contoured at 1.0*σ*. The N73 sidechain is fully enclosed by density and turns back toward the framework, where it hydrogen bonds R71 and a CDR-H2 backbone carbonyl.

**Figure S14:**
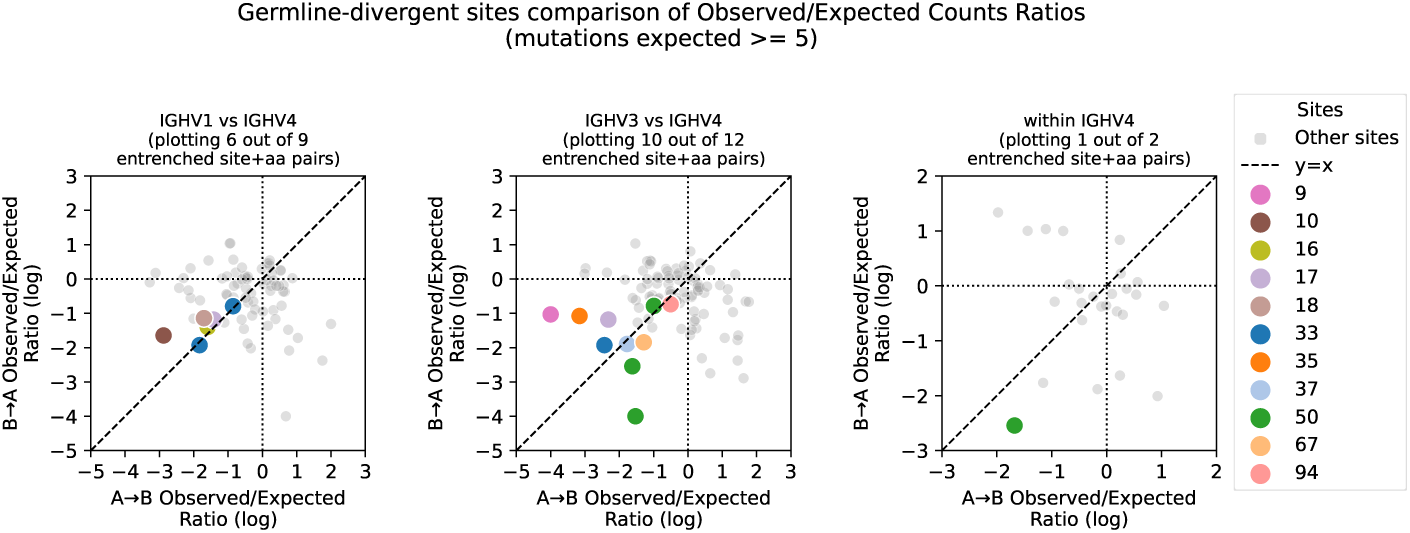
Pairwise validation of entrenched substitutions for IGHV4 comparisons. Analogous to Figure 7C but for within-IGHV4, IGHV1 vs IGHV4, and IGHV3 vs IGHV4 comparisons. Of the 23 entrenched reciprocal pairs involving IGHV4, 17 (74%) had sufficient data in both directions. All 17 showed purifying selection in both directions, with all falling below *−*0.5 and 14 (82%) below the *−*1 entrenchment threshold.

**Figure S15:**
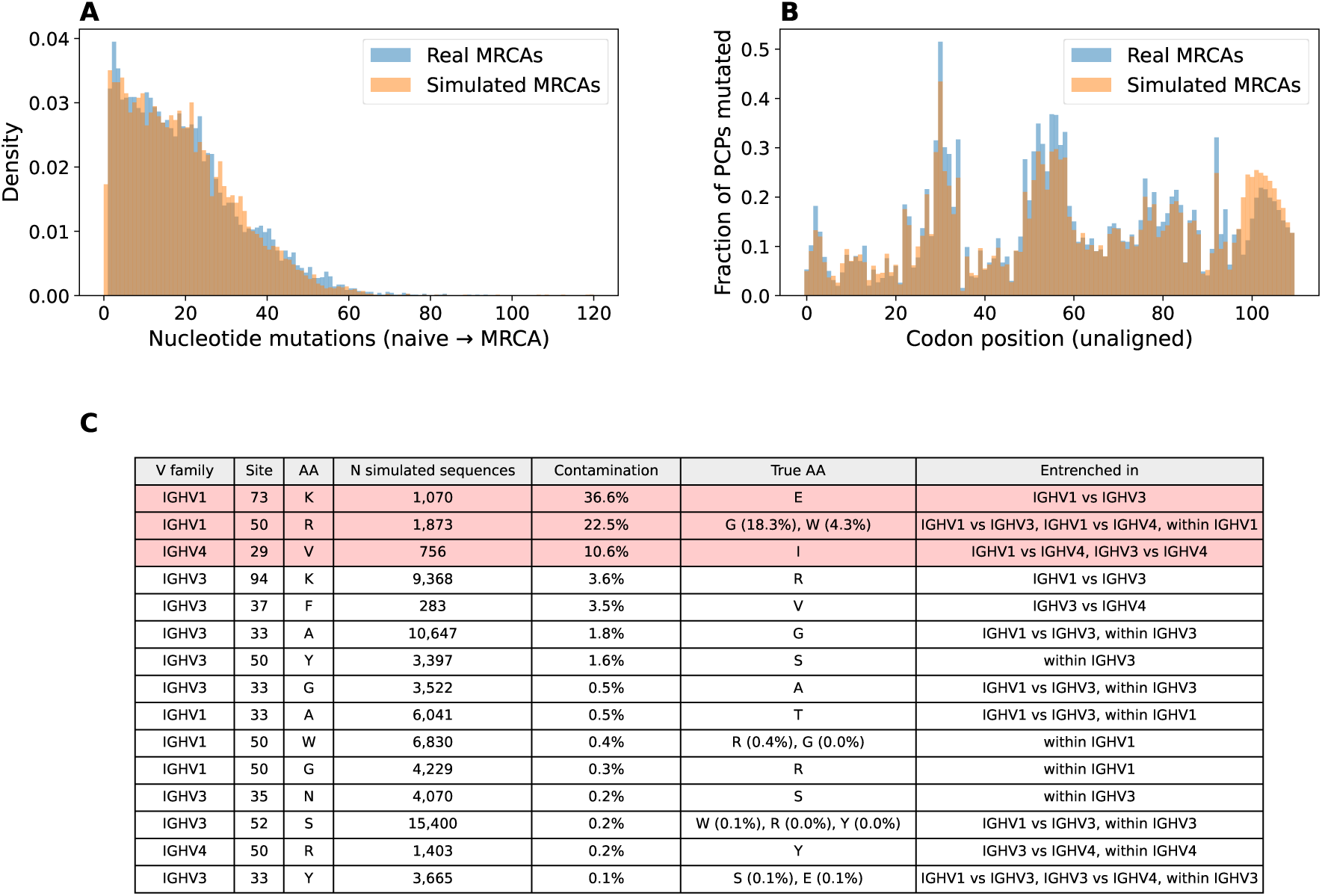
Simulations show germline allele misassignment rarely mislabels somatic mutations as germline at entrenched sites. We simulated 10 replicate clonal family root sequences per naive-to-root pair using the neutral somatic hypermutation model Thrifty and the selection model DASM, then re-annotated them with germline inference (see Methods). **(A)** Distribution of nucleotide mutation counts for real and simulated root sequences showing close agreement. **(B)** Per-codon mutation frequency along the V gene for real versus simulated root sequences. **(C)** Top 15 entrenched site/amino acid combinations ranked by contamination rate: the fraction of sequences assigned a given germline amino acid whose true germline encodes a different amino acid at that site; N is the number of simulated root sequences assigned that germline amino acid. Rows with contamination *≥*5% are highlighted in red. Three combinations have substantial contamination, all traceable to specific allele confusions (IGHV1-69, IGHV1-2, and IGHV4-59/61; see Methods); all remaining combinations have contamination below 4%, with most below 1%. At IGHV1 site 50 the entrenched amino acids other than R have negligible contamination, and at IGHV1 site 73 entrenchment holds for the K-carrying alleles that are not prone to the IGHV1-69*01/*06 confusion (Supplementary Figure S2), so neither signal rests on the contaminated group (see Methods). At IGHV4 site 29, V is observed in only one allele in our data (IGHV4-61*01), so germline-encoded entrenchment cannot be distinguished from entrenchment of somatic mutations acquired before the clonal family root.

**Figure S16:**
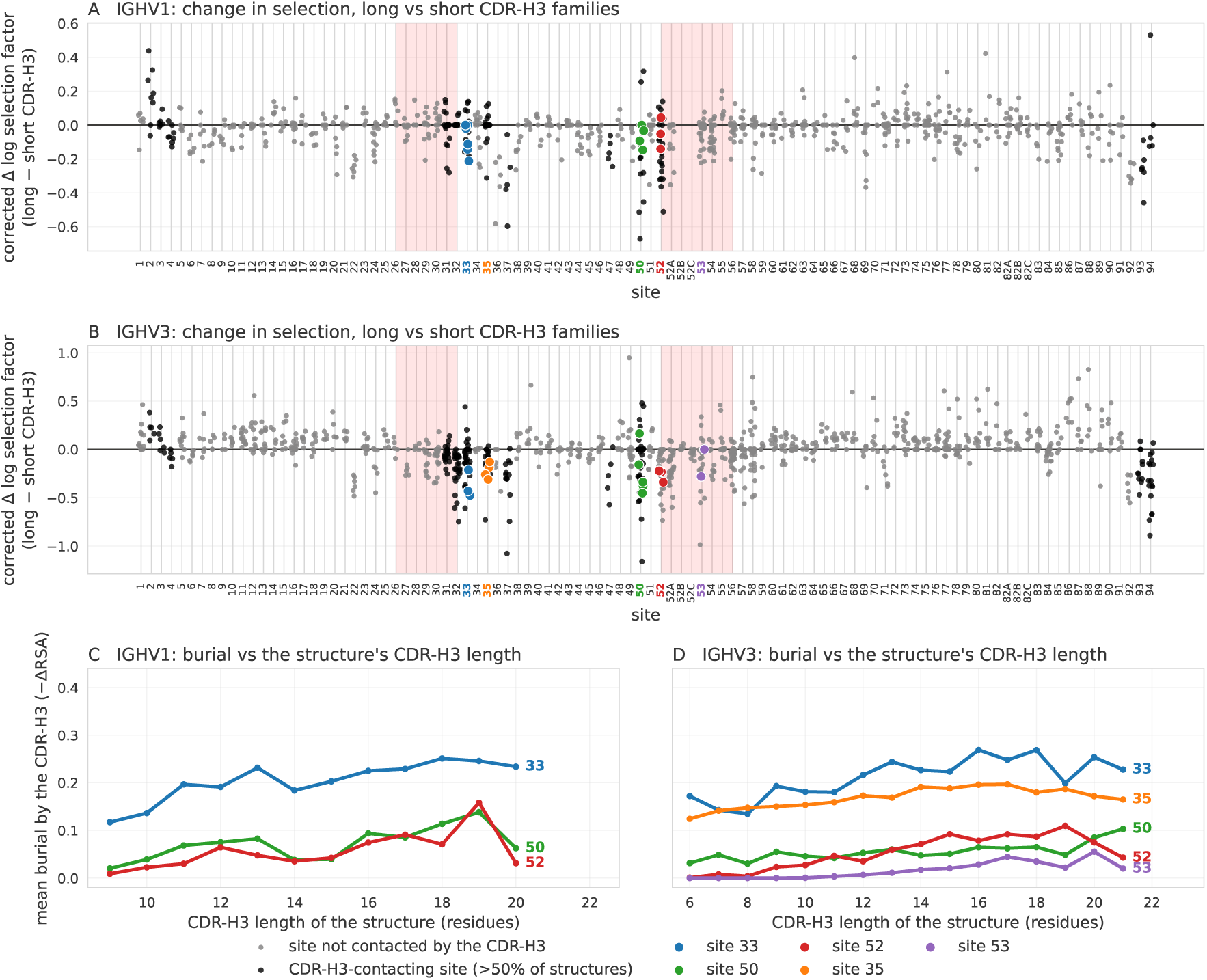
CDR-H3 length has a weak but significant effect on selection at the sites the CDR-H3 contacts, including the entrenched ones. (A, B) Each point is one substitution: the change in its median DASM log selection factor between clonal families with long and short CDR-H3s, corrected for the shift an uninformative split produces at that substitution (Methods). At the entrenched sites, colored as in previous figures, selection against leaving the germline amino acid is stronger in the families with longer CDR-H3s: 10 of 15 substitutions in IGHV1 (A) and 16 of 18 in IGHV3 (B) shift that way. The measurement is not clean, since sites elsewhere in the V region shift as well, but the largest shifts are at the sites the CDR-H3 contacts: comparing sites rather than substitutions, these move 2.7*×* as far as the rest in IGHV1 (Mann–Whitney *p* = 0.035) and 2.4*×* as far in IGHV3 (*p* = 3 *×* 10*^−^*^4^). Black marks substitutions at CDR-H3-contacting sites, grey the remainder, and pink shading highlights CDR regions. A small shift is what we expect here: a CDR-H3 has properties beyond its length that this split does not capture; the light chain and antigen vary freely within both halves of every split, adding variance that can obscure an effect; and developmental selection removes many productive IGH rearrangements before the repertoire we observe [42], so the germline–CDR-H3 combinations left to compare are only the ones that already function together. **(C, D)** Mean burial of each entrenched site by the CDR-H3 across SAbDab structures, grouped by the structure’s own CDR-H3 length. Longer CDR-H3s bury these sites more, with no model involved, which is a structural reason for the direction seen in A and B.

**Figure S17:**
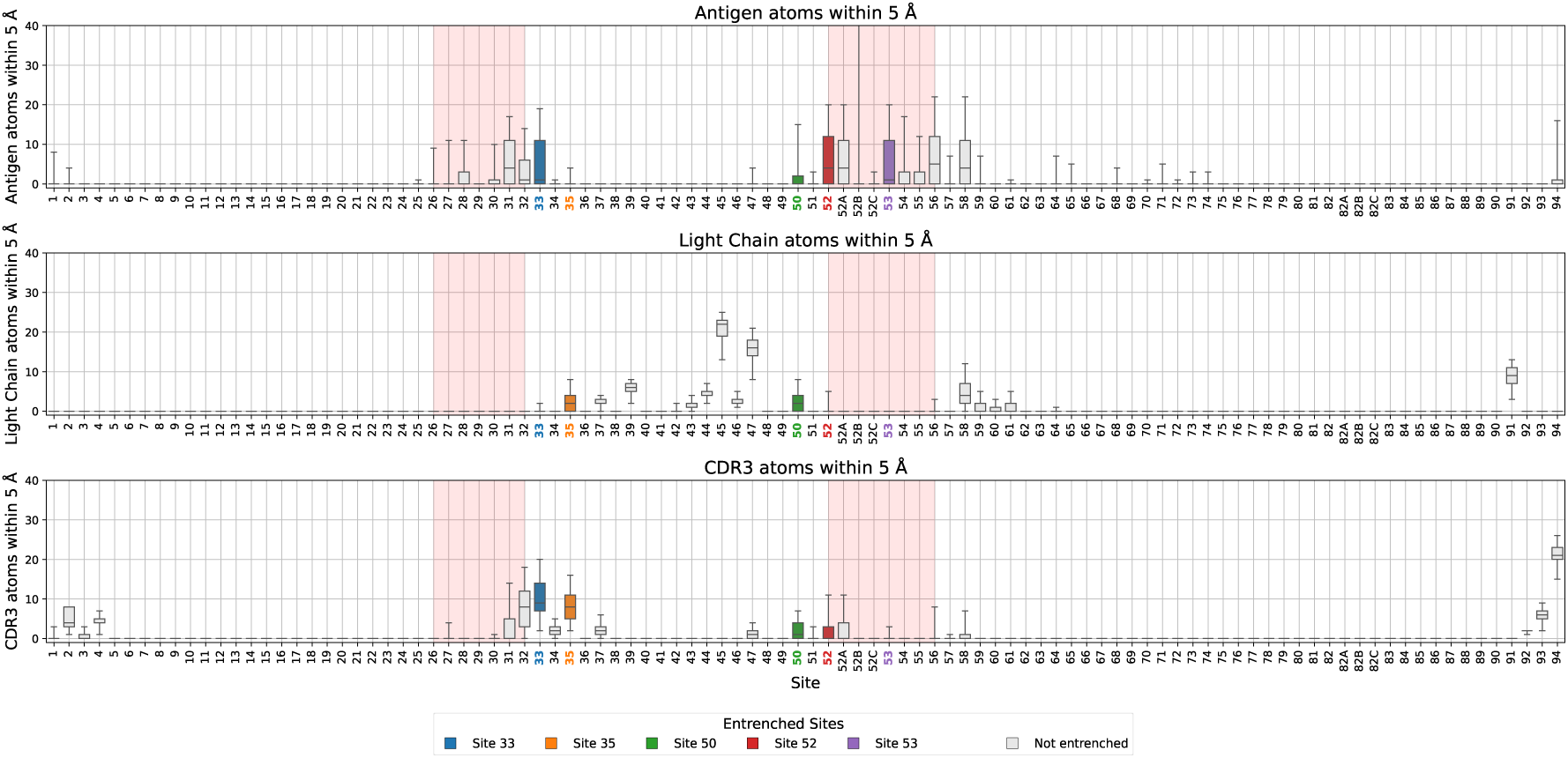
Counting atomic contacts reproduces the ΔRSA burial pattern for IGHV3. To confirm that the burial pattern in Figure 3D (rows 2–4) does not depend on measuring contact through ΔRSA, we repeated the analysis by counting atoms directly. Rows show the number of non-hydrogen atoms of the antigen, the light chain, or the CDR-H3 within 5 Å of any non-hydrogen atom of the residue. Colored boxes mark within-family entrenched sites. The distributions of contact counts show the same pattern across sites as the ΔRSA distributions in Figure 3D. Summarizing each site by a single value gives the same agreement: across sites, the fraction of structures in which a partner lies within 5 Å closely tracks the fraction in which it buries the site (ΔRSA *>* 0), with Spearman *ρ* = 0.84–0.99 for each partner in IGHV1 and IGHV3.

